# SIRT1 and SIRT3 impact host mitochondrial function and host- *Salmonella* pH balance during infection

**DOI:** 10.1101/2023.09.11.557159

**Authors:** Dipasree Hajra, Vikas Yadav, Amit Singh, Dipshikha Chakravortty

**Author notes:** Corresponding author: Prof. Dipshikha Chakravortty.

## Abstract

Mitochondria are an important organelle regulating energy homeostasis. Mitochondrial health and dynamics are crucial determinants of the outcome of several bacterial infections. SIRT3, a major mitochondrial sirtuin, along with SIRT1 regulates key mitochondrial functions. This led to considerable interest in understanding the role of SIRT1 and SIRT3 in governing mitochondrial functions during *Salmonella* infection. Here, we show that loss of SIRT1 and SIRT3 function either by shRNA-mediated knockdown or inhibitor treatment led to increased mitochondrial dysfunction with alteration in mitochondrial bioenergetics alongside increased mitochondrial superoxide generation in the *Salmonella-*infected macrophages. Consistent with dysfunctional mitochondria, mitophagy was induced along with altered mitochondrial fusion-fission dynamics in *S.* Typhimurium*-*infected macrophages. Additionally, the mitochondrial bioenergetic alteration promotes acidification of the infected macrophage cytosolic pH. This host cytosolic pH imbalance skewed the intra-phagosomal and intra- bacterial pH in the absence of SIRT1 and SIRT3, resulting in decreased SPI-2 gene expression. Our results suggest a novel role of SIRT1 and SIRT3 in maintaining the intracellular *Salmonella* niche by modulating the mitochondrial bioenergetics and dynamics in the infected macrophages.

**Graphical Abstract:** 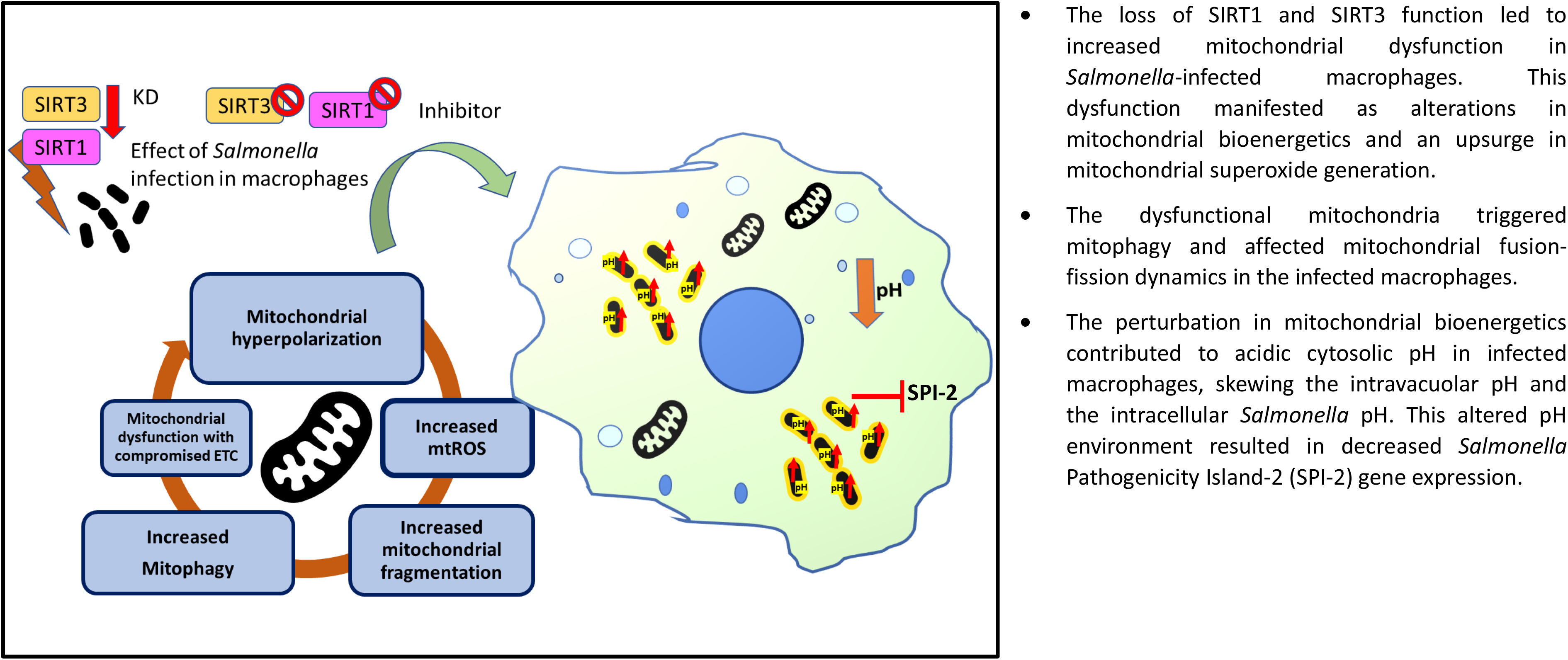

## INTRODUCTION

Mitochondria play a central role in bacterial infections, influencing host immune responses. Pathogens like *Listeria monocytogenes* and *Helicobacter pylori* induce mitochondrial fragmentation via listeriolysin O protein (LLO) (Stavru et al., 2011) or VacA effector protein (Jain et al., 2011), respectively. DRP-1- (Dynamin-related protein 1) dependent mitochondrial fragmentation is reported in both *Shigella flexneri* (Lum and Morona, 2014) and *Legionella pneumophila* infections(Escoll et al., 2017). Conversely, *Chlamydia trachomatis* infection promotes mitochondrial elongation in the early phases of infection with enhanced ATP production, facilitating *C. trachomatis* intracellular proliferation (Chowdhury et al., 2017, Kurihara et al., 2019).

Mitochondria governing TCA cycle, oxidative phosphorylation, and ATP production, are strategically targeted by *Mycobacterium tuberculosis*(Shi et al., 2015), *Brucella abortus(Czyż et al., 2017)*, and *Chlamydia trachomatis*(Siegl et al., 2014) to ensure their survival through metabolic programming.

SIRT3, a mitochondrial deacetylase, regulates mitochondrial functions including bioenergetics, metabolism, energy homeostasis (Ahn et al., 2008, Jing et al., 2011). Its role extends to ameliorating oxidative stress by inducing activation of SOD2 (superoxide dismutase 2), and CAT (catalase) that reduces Reactive oxygen species (ROS) levels(Sundaresan et al., 2009). Meanwhile, SIRT1 impact mitochondrial biogenesis via deacetylation and activation of PGC1α (Nemoto et al., 2005). Kim et al., suggested the SIRT3’s role in anti-mycobacterial host defences. Its downregulation in *Mycobacterium tuberculosis*-infected macrophages reprogramed mitochondrial metabolism and induced cell death (Smulan et al., 2021). However, in intravacuolar pathogens like *Salmonella*, the role of SIRT1 or SIRT3 in the modulation of mitochondrial function and establishment of pathogenesis is largely unexplored.

*Salmonella enterica,* causing enteric typhoid fever and gastroenteritis worldwide, intracellularly resides within macrophages, forming *Salmonella* -Containing Vacuole (SCV) after gaining into non-phagocytic intestinal epithelial cells via its *Salmonella* Pathogenicity Island-1 (SPI-1) effectors and breaching the intestinal barrier (Haraga et al., 2008). In SCV, *Salmonella* rely on SPI-2 effectors to maintain a replication niche inside their host (Haraga et al., 2008, Hajra et al., 2021).

In this study, the role of SIRT1 and SIRT3 in modulating mitochondrial bioenergetics and dynamics during *Salmonella* infection was investigated. Inhibiting SIRT1 or SIRT3 resulted in increased mitochondrial dysfunction in infected macrophages, leading to elevated mitochondrial superoxide generation and altered mitochondrial dynamics. Absence of SIRT1 or SIRT3, led to macrophage cytosol acidification, influencing intra-phagosomal and intra- bacterial pH. This shift in bacterial and phagosomal pH SPI-2 gene expression, impacting bacterial intracellular niche. Overall, SIRT1 and SIRT3 play a vital role in modulating mitochondrial bioenergetics and pH dynamics during *S*. Typhimurium infection, thereby influencing the bacterial intracellular niche and pathogenesis.

## RESULTS

### SIRT3 inhibition only led to enhanced mitochondrial superoxide generation in the infected macrophages

Mitochondrial superoxide generation is an essential arsenal of host defense and an important determinant of mitochondrial health (West et al., 2011, Suomalainen and Nunnari, 2024). *Salmonella* Typhimurium strain 14028S infection triggered an increase in mitochondrial superoxide generation in RAW264.7 murine macrophages in comparison to the uninfected control in flow cytometric studies. This increment in mitochondrial superoxide was further aggravated (around 2-fold) in knockdown condition of SIRT3 in infected RAW264.7 macrophages. However, the mitochondrial ROS production within SIRT1 knocked down macrophages remained comparable to scrambled control under infection scenario (Fig. EV1, Fig. 1 A). This result was further corroborated by the findings in the *Salmonella* infected primary peritoneal macrophages (PMs) treated with SIRT1 or SIRT3 inhibitors EX-527 (Morató et al., 2022), and 3TYP (Hu et al., 2022), respectively. Treatment with only SIRT3 inhibitor resulted in heightened production of mitochondrial ROS (mtROS) in the infected PMs in comparison to the untreated control (Fig. 1B). Immunofluorescence data further showed an increased coefficient of colocalization of mitoSOX with mitochondria in SIRT3 knockdown RAW264.7 macrophages when compared to the scrambled control (Fig. 1C-D). SIRT3 knockdown triggered increased mitochondrial oxidative stress in infected RAW264.7 macrophages, prompting the hypothesis that elevated mtROS might hinder intracellular bacterial replication. To investigate this hypothesis, we performed an intracellular bacterial proliferation assay in the presence or absence of mitochondrial superoxide scavenger, mitoTEMPO (Fig. EV2) in SIRT3 knockdown macrophages. The untreated macrophages depicted attenuated intracellular proliferation of *S.* Typhimurium in SIRT3 knockdown macrophages compared to the scrambled control. However, alleviating mtROS with mitoTEMPO, restored *Salmonella* intracellular proliferation in SIRT3 knockdown macrophages (Fig.1E). To ensure that the observed reduction in intracellular burden in the shSIRT3 macrophages was not due to increased cell death, a cell death assay was performed (Fig. EV3). Results suggested no defect in the cell viability upon SIRT3 knockdown both in the untreated and infected macrophages. Therefore, the increased mitochondrial oxidative burst within SIRT3 knockdown macrophages limits *Salmonella* intracellular proliferation, a restriction reversed upon alleviating mtROS production by mitoTEMPO.

**Fig. 1.**
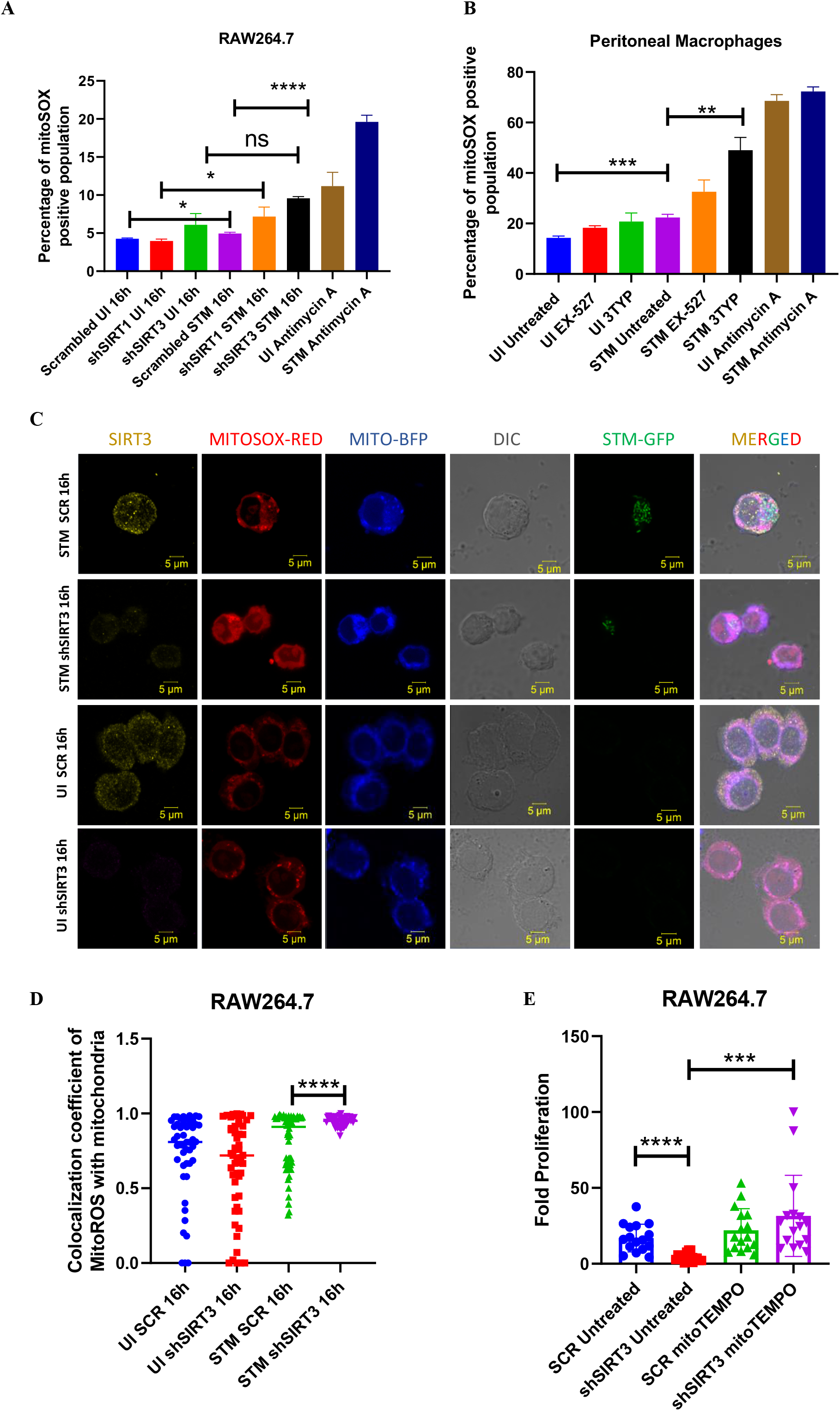
SIRT3 inhibition leads to enhanced mitochondrial ROS generation in infected macrophages leading to attenuated *Salmonella* replication. A-Mitochondrial ROS estimation via flow cytometry in *S.* Typhimurium infected RAW264.7 macrophages at 16hr post infection under SIRT1 or SIRT3 knockdown condition. Data is representative of N=4, n=2. 10µM antimycin A-treated (ROS inducer) uninfected and infected cells served as positive control. Unpaired two-tailed Student’s t test was performed to obtain the p values. *** p < 0.001, ** p<0.01, * p<0.05 B-Mitochondrial ROS estimation via flow cytometry in *S.* Typhimurium infected peritoneal macrophages (isolated post 5^th^ day of thioglycolate injection) at 16hr post infection in the presence of SIRT1 (EX-527,1µM) or SIRT3 (3TYP, 1µM) inhibitor treatment. Data is representative of N=3, n=2. Unpaired two-tailed Student’s t test was performed to obtain the p values. *** p < 0.001, ** p<0.01, * p<0.05 C-D- Representative confocal images of SIRT3 knockdown RAW264.7 macrophages exhibiting co-localization coefficient of mitoSox Red detecting mitochondrial ROS with mitochondria (mitoBFP) upon *S.* Typhimurium infection at indicated time points post-infection. Data is representative of N=3, n>50 (microscopic field). Unpaired two-tailed Student’s t test was performed to obtain the p values. *** p < 0.001, ** p<0.01, * p<0.05 E- Intracellular proliferation assay of *S*. Typhimurium within scrambled control or SIRT3 knockdown macrophages in presence or absence of mitoTEMPO treatment. Data is representative of N=3, n=2. Unpaired two-tailed Student’s t test was performed to obtain the p values. *** p < 0.001, ** p<0.01, * p<0.05

Together, these results implicate the role of SIRT3 in mitigating mitochondrial ROS generation in the *Salmonella-*infected murine macrophages and thereby facilitating bacterial proliferation.

### SIRT1 and SIRT3 inhibition compromised the mitochondrial antioxidant defence in the *S.* Typhimurium-infected macrophages

Our prior data suggested increased mitochondrial ROS-mediated mitochondrial oxidative stress in SIRT3 knockdown RAW264.7 macrophages in *S*. Typhimurium infected macrophages. Considering the host’s mitochondrial antioxidant defense mechanisms crucial in countering oxidative stress (Forman and Zhang, 2021, Chen et al., 2018), we conducted RT-qPCR studies of several mitochondrial antioxidant genes (*Gpx*, *Ho-1*, *Cat*, and *Sod2*) upon SIRT1 and SIRT3 knockdown in RAW264.7 macrophages and in peritoneal macrophages treated with SIRT1 (EX-527) or SIRT3 (3TYP) inhibitors. Both SIRT1 and SIRT3 knockdown, as well as inhibitor treatments, resulted in decreased transcript levels of mitochondrial antioxidant genes in both RAW264.7 and peritoneal macrophages during infection (Fig. 2A-B). Subsequently, we evaluated the status of bacterial antioxidant gene expression within SIRT1 or SIRT3 knockdown macrophages. In shSIRT3 macrophages, there was a notable decrease in bacterial antioxidant gene expression (*sodA*, *sodB*, *katE*, *katG*) compared to the scrambled control. Conversely, shSIRT1-infected macrophages depicted heightened bacterial antioxidant gene expression compared to the scrambled control. (Fig. EV4). Previous literature implicates SIRT3 deacetylating SOD2 at critical lysine residues (K53 and K89), promoting its antioxidant activity(Qiu et al., 2010). To validate this interaction during *S.* Typhimurium infection condition, immunoprecipitation studies were performed. Immunoprecipitation studies revealed the interaction of SOD2 with SIRT3 upon infection. However, this interaction got dampened in SIRT3 knockdown, and inhibitor-treated infected RAW264.7 macrophages, alongside SOD2 hyperacetylation (Fig. 2C-D). Further, SOD2 activity assay demonstrated reduced active SOD2 or MnSOD2 concentration in the *S*. Typhimurium-infected RAW264.7 macrophages under both SIRT3 knockdown or inhibited conditions (Fig. 2E-F). Together, our results highlight the role of SIRT1 and more predominantly SIRT3 in rendering protection against mitochondrial oxidative stress upon *Salmonella* infection by deacetylating SOD2 and thereby promoting its antioxidant properties.

**Fig. 2.**
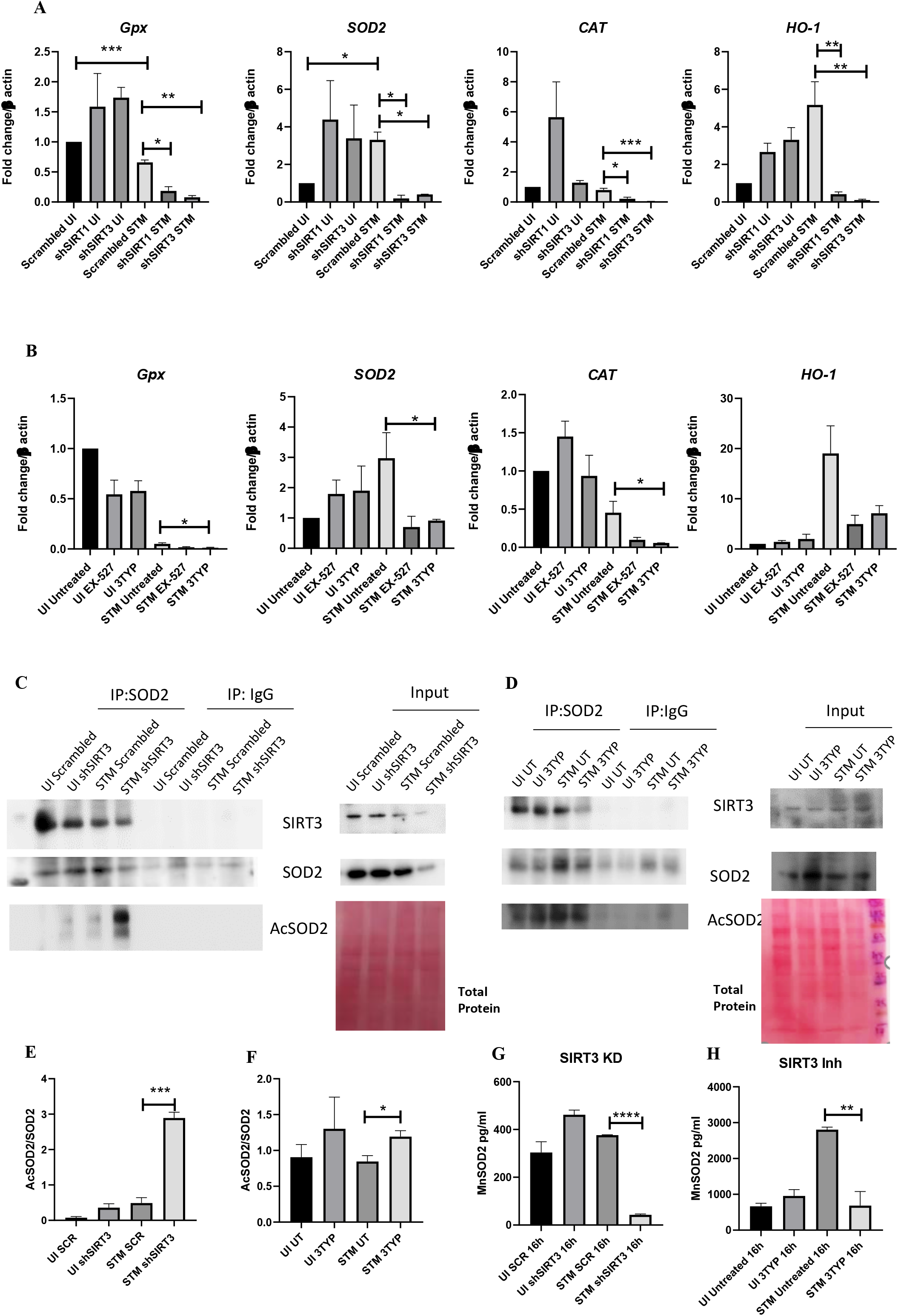
Effect of SIRT1 or SIRT3 inhibition on host mitochondrial antioxidant defense in *Salmonella* Typhimurium infection scenario. A-Quantitative gene expression studies of several host mitochondrial antioxidant genes such as *Gpx*, *Sod2*, *Cat*, and *Ho-1* in RAW264.7 macrophages upon knockdown of SIRT1 and SIRT3 in *S.* Typhimurium infection scenario. Data is representative of N=3, n=2. Unpaired two-tailed Student’s t test was performed to obtain the p values. *** p < 0.001, ** p<0.01, * p<0.05 B-Quantitative gene expression studies of several host mitochondrial antioxidant genes such as *Gpx*, *Sod2*, *Cat*, and *Ho-1* in peritoneal macrophages (isolated from 6–8-week- old adult male mice post 5^th^ day of thioglycolate injection) after 16h of *S.* Typhimurium infection upon SIRT1 (EX-527,1µM) or SIRT3 (3TYP, 1µM) inhibitor treatment. Data is representative of N=3, n=2. Unpaired two-tailed Student’s t test was performed to obtain the p values. *** p < 0.001, ** p<0.01, * p<0.05 C- Immunoprecipitation studies depicting SOD2 interaction with SIRT3 upon immunoprecipitation of SOD2 in uninfected (UI) or *S.* Typhimurium (STM) infected SIRT3 knockdown RAW264.7 macrophages at 16hr post-infection. Data is representative of N=2, n=1. *** p < 0.001, ** p<0.01, * p<0.05 D- Immunoprecipitation studies depicting SOD2 interaction with SIRT3 upon immunoprecipitation of SOD2 in uninfected (UI) or *S.* Typhimurium (STM) infected RAW264.7 macrophages at 16hr post-infection upon inhibitor treatment of SIRT3. Data is representative of N=2, n=1. *** p < 0.001, ** p<0.01, * p<0.05 E- Mitochondrial SOD2 activity assay in uninfected and *S*. Typhimurium infected RAW264.7 macrophages in the knockdown condition of SIRT3. Data is representative of N=2, n=2. Unpaired two-tailed Student’s t test was performed to obtain the p values. *** p < 0.001, ** p<0.01, * p<0.05 F- Mitochondrial SOD2 activity assay in uninfected and *S*. Typhimurium infected RAW264.7 macrophages upon SIRT3 inhibitor treatment. Data is representative of N=2, n=2. Unpaired two-tailed Student’s t test was performed to obtain the p values. *** p < 0.001, ** p<0.01, * p<0.05

### SIRT1 or SIRT3 inhibition resulted in compromised ETC function and lowered ATP generation in the infected macrophages

SIRT3, crucial for maintaining healthy mitochondrial homeostasis, is associated with ATP synthase subunit (Yang et al., 2016) and regulates basal ATP levels in vivo (Ahn et al., 2008). Therefore, to explore the impact of SIRT3 inhibition or downregulation on ATP production during *Salmonella* infection, an ATP estimation assay was performed. *Salmonella*-infected SIRT1 or SIRT3 RAW264.7 macrophages exhibited decreased ATP production in the later phases of infection at 16hr post-infection compared to the infected scrambled or un-transfected control (Fig. 3A). Moreover, earlier data showed exacerbated production of mtROS upon SIRT3 knockdown or inhibition in *Salmonella*-infected murine macrophages. Given that both mtROS production and ATP synthesis are governed by mitochondrial ETC (Electron transport chain), we investigated whether SIRT1 and SIRT3 inhibition or knockdown disrupts ETC function during *Salmonella* infection. qPCR-mediated ETC Complex gene profiling in infected RAW264.7 macrophages under SIRT1 or SIRT3 knockdown conditions revealed decreased expression of Complex I subunit genes (*NDUFV1*, *NDUFS1*, *NDUFA9*), Complex II (*SDHD),* Complex III (*UQCR*), Complex IV (*COX5B*) and Complex V genes (*ATP5B*) (Fig. 3B). Similar findings were obtained from infected peritoneal macrophages treated with SIRT1 or SIRT3 catalytic inhibitors (Fig. 3C). Consequently, SIRT1 or SIRT3 knockdown or inhibition triggers mitochondrial ETC dysfunction in infected macrophages, potentially leading to enhanced mitochondrial ROS generation and reduced ATP production.

**Fig. 3.**
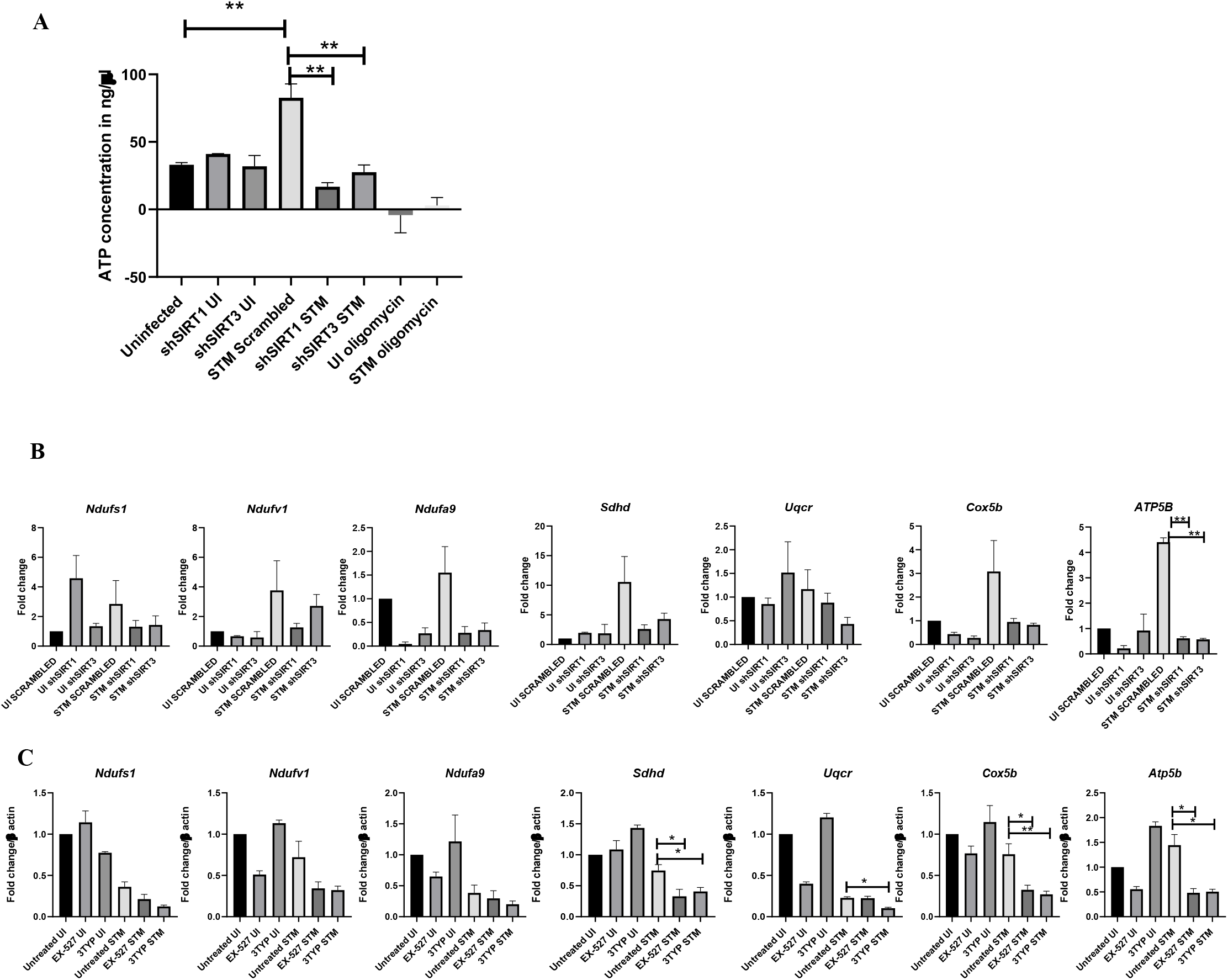
Inhibition of SIRT1 or SIRT3 leads to disruption of mitochondrial bioenergetics in *S.* Typhimurium-infected macrophages. A- ATP estimation assay in SIRT1 or SIRT3 knockdown RAW264.7 macrophages in the presence or absence of *S.* Typhimurium infection at specified time points post-infection. Data is representative of N=3, n=2. 10µM oligomycin-treated (ATP synthase inhibitor) uninfected and infected cells served as negative control. Unpaired two-tailed Student’s t test was performed to obtain the p values. *** p < 0.001, ** p<0.01, * p<0.05 B- qRTPCR mediated expression analysis of several Complex I, Complex II, Complex III, Complex IV, and Complex V genes of the Electron Transport Chain in *S.* Typhimurium infected RAW264.7 macrophages at 16hr post infection under the knockdown condition of either SIRT1 or SIRT3. Data is representative of N=3, n=2. Unpaired two-tailed Student’s t test was performed to obtain the p values. *** p < 0.001, ** p<0.01, * p<0.05 C- qRTPCR mediated expression analysis of several Complex I, Complex II, Complex III, Complex IV, and Complex V genes of the Electron Transport Chain in *S.* Typhimurium infected peritoneal macrophages at 16hr post-infection in the presence or absence of SIRT1 (EX-527) or SIRT3 (3TYP) inhibitor treatment at a concentration of 1µM. Data is representative of N=3, n=2. Unpaired two-tailed Student’s t test was performed to obtain the p values. *** p < 0.001, ** p<0.01, * p<0.05

### SIRT3 modulated the mitochondrial membrane potential of the infected macrophages

Mitochondrial function, ATP production and mtROS generation rely on mitochondrial membrane potential. To investigate alteration in the mitochondrial membrane potential during *S.* Typhimurium infection, we undertook TMRM staining of the infected macrophages in SIRT1 or SIRT3 knockdown condition through flow cytometry. TMRM (Tetramethylrhodamine Methyl Ester, Perchlorate) is a cell-permeant fluorescent dye assessing mitochondrial membrane potential (Chen et al., 2002). TMRM staining of the infected RAW 264.7 macrophages revealed enhanced membrane potential under the knockdown condition of SIRT3 only at 16hr post-infection compared to the scrambled control (Fig. 4). SIRT1 knockdown alone did not exert significant alterations in mitochondrial membrane potential within infected macrophages (Fig. 5). Mitochondrial hyperpolarization has been shown to be caused by Complex I inhibition which subsequently triggers reduced activity of Complex II, III, and IV and reduced forward action of Complex V(Forkink et al., 2014). Our data of mitochondrial hyperpolarization in *Salmonella*-infected RAW264.7 macrophages coincides with decreased expression of Complex I, II, III, IV and V (as shown earlier).

**Fig. 4.**
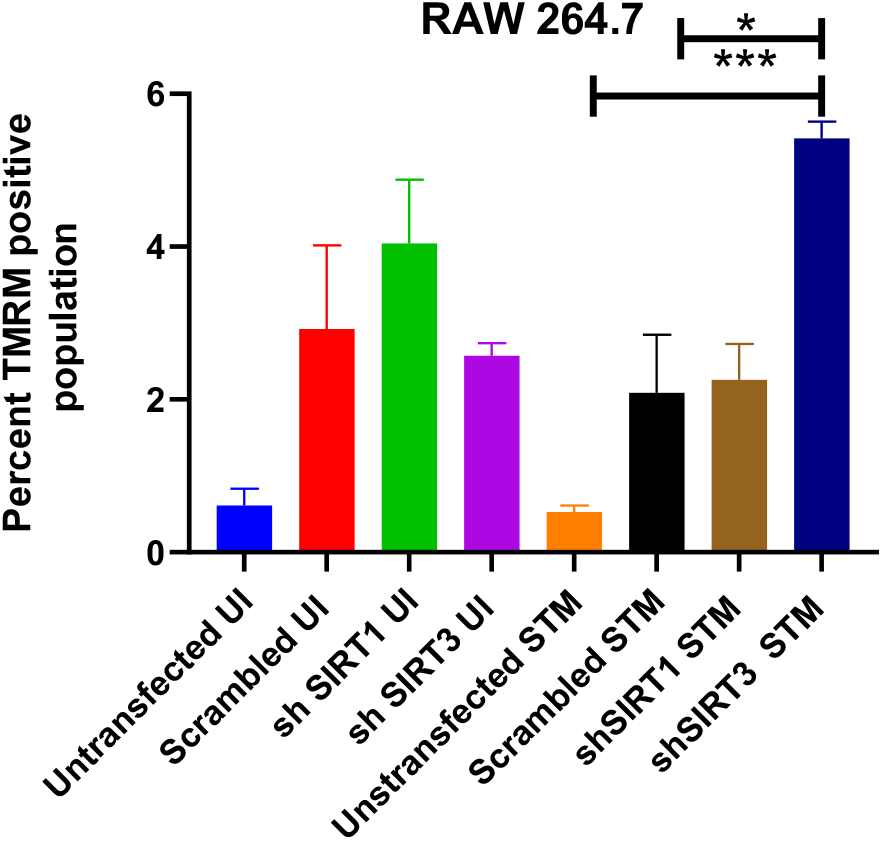
SIRT3 inhibition mediates mitochondrial membrane hyperpolarization. Mitochondrial membrane potential determination using TMRM dye via flow cytometry in RAW264.7 macrophages under knockdown condition of SIRT1 or SIRT3 at 16hr post infection. Data is representative of N=3, n=2. Unpaired two-tailed Student’s t test was performed to obtain the p values. *** p < 0.001, ** p<0.01, * p<0.05

**Fig. 5.**
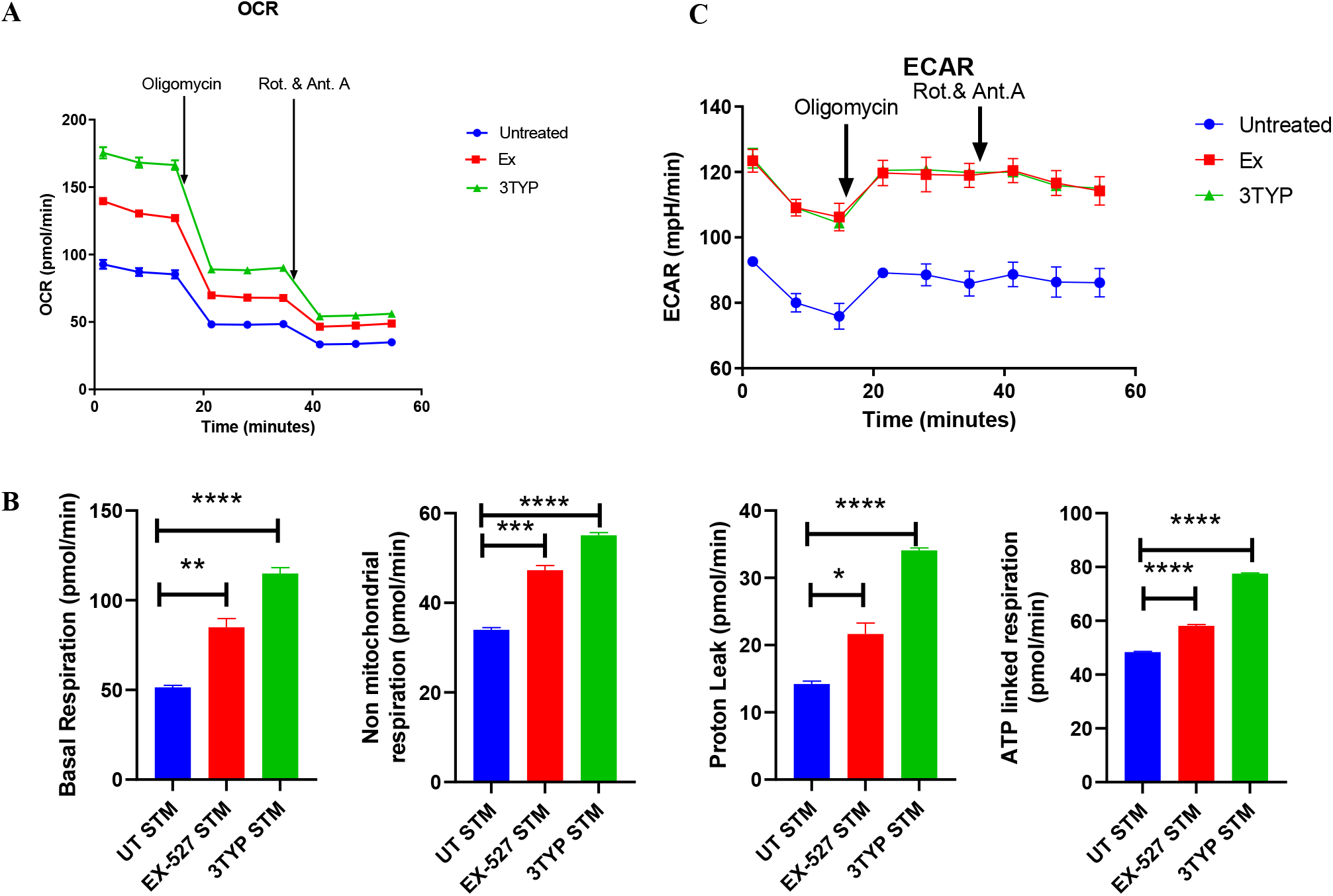
Mito stress test of *S.* Typhimurium infected peritoneal macrophages under SIRT3 inhibitor treatment. A-Mitochondrial Oxygen Consumption Rate (OCR) curve of infected peritoneal macrophages isolated from 6-8 week old adult male mice post 5^th^ day of thioglycolate injection in the presence or absence of SIRT1 (EX-527) or SIRT3 inhibitor (3TYP,1µM) treatment. Data is representative of N=3, n=2. Unpaired two-tailed Student’s t test was performed to obtain the p values. *** p < 0.001, ** p<0.01, * p<0.05 B- Basal mitochondrial respiration of infected peritoneal macrophages isolated from 6– 8-week-old adult male mice post 5^th^ day of thioglycolate injection in the presence or absence of SIRT3 inhibitor (3TYP,1µM) treatment as calculated from A. Data is representative of N=3, n=2. Unpaired two-tailed Student’s t test was performed to obtain the p values. *** p < 0.001, ** p<0.01, * p<0.05 C- Non-mitochondrial respiration of infected peritoneal macrophages upon SIRT1 and SIRT3 inhibitor treatment post 6h of *S*. Typhimurium infection as shown in A. Data is representative of N=3, n=2. Unpaired two-tailed Student’s t test was performed to obtain the p values. *** p < 0.001, ** p<0.01, * p<0.05 D- Proton leak of infected peritoneal macrophages upon SIRT1 and SIRT3 inhibitor treatment post 6h of *S*. Typhimurium infection as obtained from A. Data is representative of N=3, n=2. Unpaired two-tailed Student’s t test was performed to obtain the p values. *** p < 0.001, ** p<0.01, * p<0.05 E-ATP-linked respiration infected peritoneal macrophages upon SIRT1 and SIRT3 inhibitor treatment post 6h of *S*. Typhimurium infection as obtained from A. Data is representative of N=3, n=2. Unpaired two-tailed Student’s t test was performed to obtain the p values. *** p < 0.001, ** p<0.01, * p<0.05 F-Extracellular Acidification Rate (ECAR) profile of infected peritoneal macrophages upon SIRT1 and SIRT3 inhibitor treatment post 6h of *S*. Typhimurium infection. Data is representative of N=3, n=2. Unpaired two-tailed Student’s t test was performed to obtain the p values. *** p < 0.001, ** p<0.01, * p<0.05

### SIRT1 and SIRT3 inhibition led to altered mitochondrial respiration in *S*. Typhimurium infected macrophages

Upon observing reduced ATP production following SIRT1 or SIRT3 inhibition, we determined mitochondrial bioenergetics in infected peritoneal macrophages under SIRT1 and SIRT3 inhibitor treatment. Using a modified Seahorse Mito stress test (Cumming et al., 2018), we assessed the mitochondrial oxygen consumption rates (OCR) post 6hr of infection and inhibitor treatment in peritoneal macrophages over a 60 min interval (Fig. 5A). The baseline OCR provided basal respiration by eliminating the non-mitochondrial respiration. Oligomycin addition allowed computation of ATP-linked OCR and proton leakage, while Antimycin A and rotenone revealed the rate of non-mitochondrial respiration. Simultaneously, the extracellular acidification rate (ECAR), which is proportional to the lactate efflux, was measured and interpreted as an indicator of glycolytic flux in the macrophages. Results revealed increased basal respiration, non-mitochondrial respiration, proton leakage, and mitochondrial ATP-linked respiration (Fig. 5B) and an increase in ECAR in the infected peritoneal macrophages under both SIRT1 (EX-527) and SIRT3 (3TYP) catalytic inhibitor treatment compared to the infected untreated control (Fig. 5C). This underscores the function of SIRT1 and SIRT3 in regulating mitochondrial bioenergetics during *Salmonella* infection (Fig. 5). The overall increase in mitochondrial respiratory flux and proton leakage may contribute to increased mitochondrial ROS production and mitochondrial hyperpolarization (Amorim et al., 2022) (as depicted earlier). Previous reports suggest enhanced respiration alongside decreased ATP production may involve compensation through increased glycolysis (Hajra et al., 2022) or non-mitochondrial respiration (Yang et al., 2021). Additionally, heightened respiration leading to increased superoxide generation can result in proton leakage across the inner mitochondrial membrane and decreased ATP synthesis (BUTLER et al., 2003, Amorim et al., 2022).

### pH homoeostasis within *S.* Typhimurium infected macrophages is mediated by SIRT3

Our previous findings showed defects in mitochondrial bioenergetics, ETC function loss, increased mtROS generation, mitochondrial membrane hyperpolarization and lowered ATP production in STM-infected macrophages upon SIRT3 or SIRT1 inhibition. Considering the potential impact of mitochondrial membrane potential alterations on cytosolic pH(Bonora et al., 2022, Poburko et al., 2011), we examined the cytosolic pH of STM-infected RAW264.7 macrophages under the knockdown condition of SIRT1 and SIRT3 by BCECF-AM staining. Host cytosolic pH sensing is crucial for *Salmonella* intravacuolar life, governing effector translocation (Yu et al., 2010). BCECF-AM, a cell-permeant dye, measures pH-dependent emission intensity ratios at 535nm with dual excitation at 488nm and 405nm. Knockdown of SIRT3, not SIRT1, within *S.* Typhimurium-infected RAW264.7 macrophages intensified the cytosolic acidity of the macrophages (pH∼4.8) (Fig.6A-B). Similar observations were obtained from the peritoneal macrophages wherein SIRT3 enzymatic inhibitor treatment resulted in decreased fluorometric ratio of BCECF-AM, indicative of lowered intracellular pH (pH∼4.43) (Fig.6C-D). Altogether, our findings suggest the role of SIRT3 in maintaining cytosolic pH through regulation of mitochondrial membrane potential and bioenergetics.

**Fig. 6.**
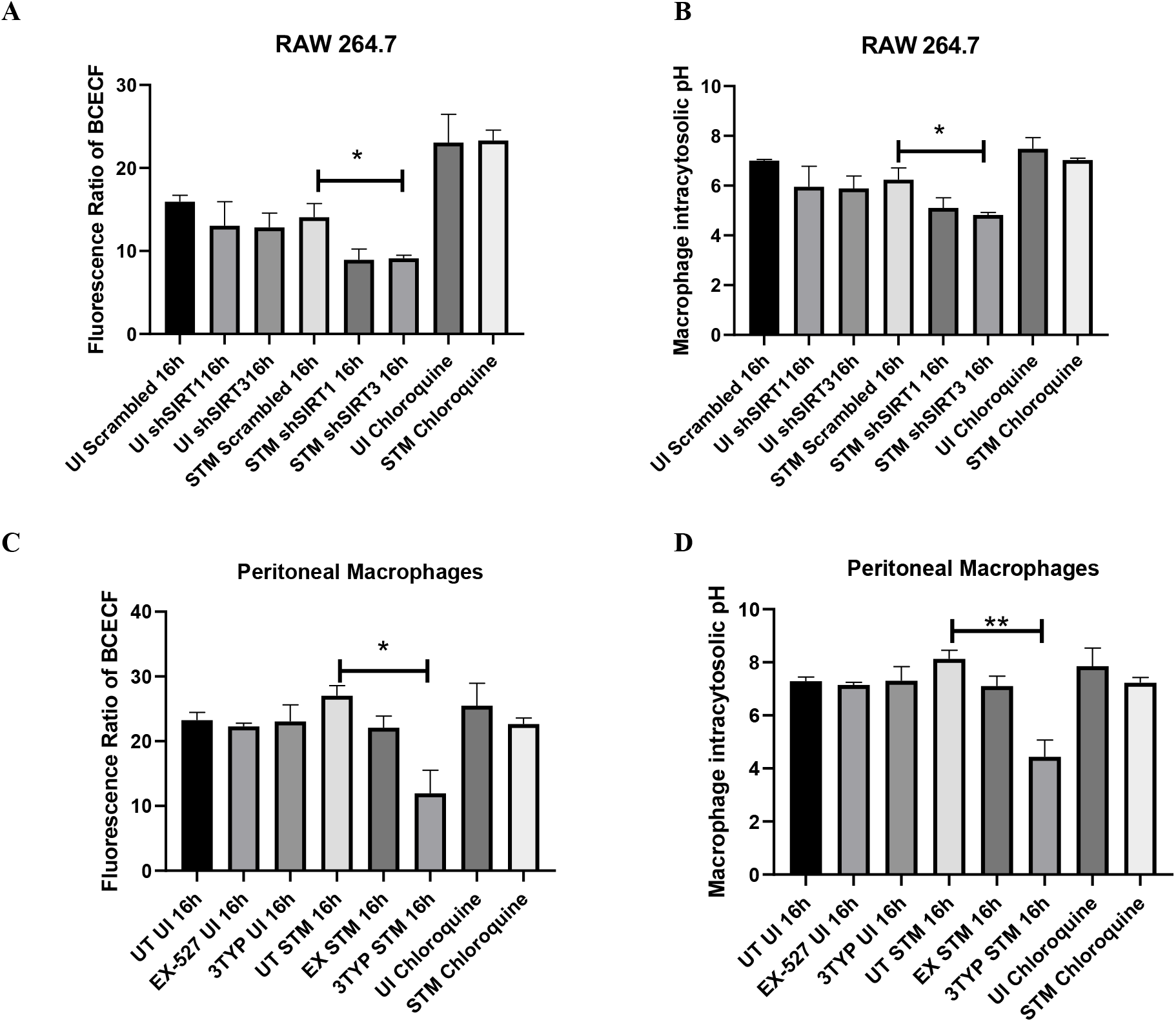
SIRT3 knockdown causes alteration in the intracellular cytosolic pH of the infected macrophages. A-B-Flow cytometric analysis of cytosolic pH of *S.* Typhimurium infected RAW264.7 macrophages using BCECF-AM dye under knockdown conditions of SIRT1 and SIRT3 or 50µg/ml of chloroquine treatment. Data is representative of N=4, n=3. Unpaired two-tailed Student’s t test was performed to obtain the p values. *** p < 0.001, ** p<0.01, * p<0.05 C-D-Flow cytometric analysis of cytosolic pH of *S.* Typhimurium infected peritoneal macrophages using BCECF-AM dye under inhibitor treatment of SIRT1 (EX-527) and SIRT3 (3TYP) inhibitor at the concentration of 1µM or 50µg/ml of chloroquine treatment. Data is representative of N=4, n=3. Unpaired two-tailed Student’s t test was performed to obtain the p values. *** p < 0.001, ** p<0.01, * p<0.05

### SIRT3 triggered host cytosolic pH alterations influenced both intra-phagosomal and intra-*Salmonella* pH and its T3SS mediated virulence protein secretion

Previous reports suggested the prerequisite for host cytosol sensing by *Salmonella* to facilitate *Salmonella* Pathogenicity Island 2 (SPI-2) (Chakravortty et al., 2005) encoded complex dissociation, degradation, and effector secretion(Yu et al., 2010). In light of this, we hypothesized that alterations in host cytosolic pH alteration due to SIRT1 or SIRT3 knockdown or inhibition might influence intra-phagosomal and intra-bacterial pH, subsequently impacting SPI-2 effector secretion. Examination of SCV pH within the pH rhodo-labelled STM-infected SIRT1 or SIRT3 RAW264.7 macrophages revealed an increase in intra-vacuolar pH (pH∼7.0) when compared to the scrambled control (pH∼5.7 to 6.0) (Fig. 7A, Fig. EV5A). Similarly, SIRT1 or SIRT3 inhibitor-treated RAW264.7 macrophages exhibited a loss in intra-phagosomal acidification (pH∼7.0) relative to the untreated control (pH∼4.5 to 5.8) (Fig. 7B, Fig. EV5B). Subsequently, intra-bacterial pH within the SIRT1 and SIRT3 knockdown macrophages were evaluated by infecting RAW264.7 macrophages with *S.* Typhimurium 14028S expressing pBAD-pHuji (Shen et al., 2014) (a plasmid encoding pH-sensitive red fluorescent protein). The intra-*Salmonella* pH in SIRT1 or SIRT3 knockdown macrophages exhibited a loss in acidic pH, approaching neutrality (pH∼7.2 to 7.6) compared to the scrambled control (pH∼5.4) (Fig.7C). Similar results were obtained in inhibitor-treated peritoneal macrophages, wherein both SIRT1 and SIRT3 inhibitor treatment resulted in a loss of acidification in the intra-bacterial pH (Fig.7D). In addition to host cytosolic pH, SPI-2 effector secretion is influenced by the acidic pH of the bacterial cytosol, activating OmpR, a regulator of SPI-2(Chakraborty et al., 2015). Inside the macrophages, the cadaverine operon *cadC/B* was repressed by OmpR and the *Salmonella* cytoplasm remained acidified. In *ompR* deficient *Salmonella,* the intracellular pH is returned to a near neutral in a CadC/BA-dependent manner. This activation of EnvZ/OmpR by acid stress stimulates the production of the SsrA/B, another two-component system, and subsequently, expression of SPI-2-secreted effectors(Chakraborty et al., 2015, Garmendia et al., 2003, Wang et al., 2012). This phenomenon holds true in bacteria residing inside the SIRT1 and SIRT3 knockdown macrophages wherein the *S*. Typhimurium within the knockdown or inhibitor treated macrophages exhibit increased pH like that of *ompR* deficient *S*. Typhimurium strain and consequently exhibit increased expression of *cadA*, *cadB* and *cadC* together amounting to decreased SPI-2 gene expression such as *pipB2*, and *steE* (Fig.7E-I).

**Fig. 7.**
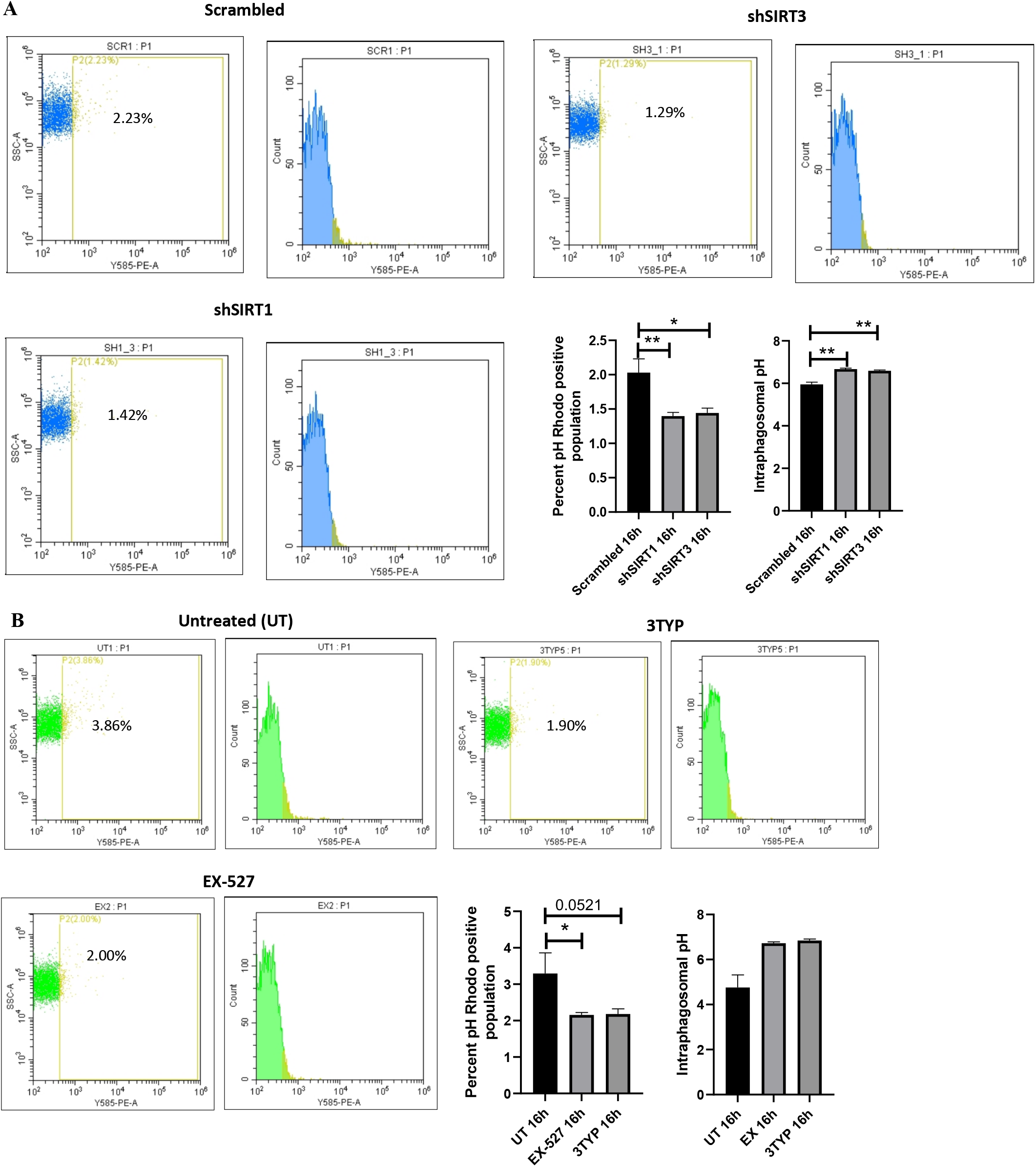
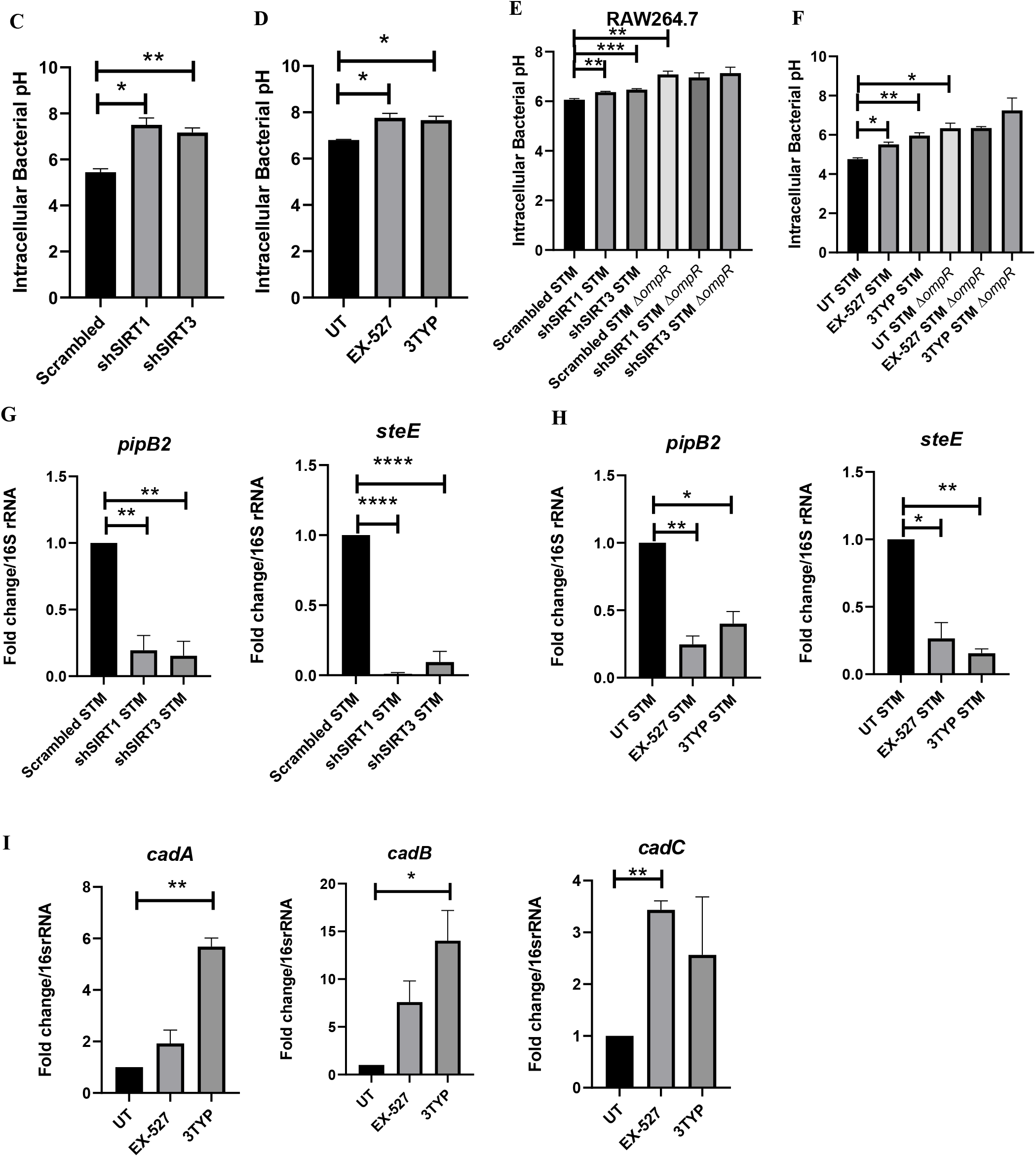
SIRT1 and SIRT3 knockdown or inhibition trigger loss in acidification of bacterial pH and subsequently its SPI-2 secretion within infected macrophages. A-B- Flow cytometric analysis of SCV pH of infected RAW264.7 macrophages under knockdown condition of SIRT1 or SIRT3 at 16h post-infection. Data is representative of N=2,n=3. Unpaired two-tailed Student’s t-test was performed to obtain the p-values. *** p < 0.001, ** p<0.01, * p<0.05 C-D- Flow cytometric analysis of SCV pH of infected RAW264.7 macrophages under inhibitor treatment of SIRT1 or SIRT3 at 16h post-infection. Data is representative of N=2,n=3. Unpaired two-tailed Student’s t-test was performed to obtain the p-values. *** p < 0.001, ** p<0.01, * p<0.05 G- Intra-bacterial pH of *S.* Typhimurium expressing pH-sensitive plasmid pHUji was measured from infected RAW264.7 macrophages using flow cytometry under knockdown conditions of SIRT1 and SIRT3. Data is representative of N=3, n=4. Unpaired two-tailed Student’s t-test was performed to obtain the p values. *** p < 0.001, ** p<0.01, * p<0.05 H-Intra-bacterial pH of *S.* Typhimurium expressing pH-sensitive plasmid pHUji was measured from infected peritoneal macrophages using flow cytometry under inhibitor treatment of SIRT1 (EX-527) and SIRT3 (3TYP) inhibitor at the concentration of 1µM. Data is representative of N=3, n=2. Unpaired two-tailed Student’s t-test was performed to obtain the p values. *** p < 0.001, ** p<0.01, * p<0.05 I-Intra-bacterial pH of *S.* Typhimurium WT (STM WT) and STMΔ pH-sensitive plasmid pHUji was measured from infected RAW264.7 macrophages using flow cytometry under knockdown conditions of SIRT1 and SIRT3. Data is representative of N=3, n=4. Unpaired two-tailed Student’s t-test was performed to obtain the p values. *** p < 0.001, ** p<0.01, * p<0.05 J- Intra-bacterial pH of *S.* Typhimurium WT (STM WT) and STMΔ pH-sensitive plasmid pHUji was measured from infected RAW264.7 macrophages using flow cytometry under inhibitor treatment of SIRT1 (EX-527) and SIRT3 (3TYP) inhibitor at the concentration of 1µM. Data is representative of N=3, n=4. Unpaired two-tailed Student’s t-test was performed to obtain the p values. *** p < 0.001, ** p<0.01, * p<0.05 K- Quantitative PCR-mediated expression studies of SPI-2 genes within infected RAW264.7 macrophages under the knockdown condition of SIRT1 and SIRT3. Data is representative of N=3, n=4. Unpaired two-tailed Student’s t-test was performed to obtain the p values. *** p < 0.001, ** p<0.01, * p<0.05 L- Quantitative PCR-mediated expression studies of SPI-2 genes within infected peritoneal macrophages under the inhibitor treatment of SIRT1 (EX-527) and SIRT3 (3TYP) inhibitors. Data is representative of N=3, n=4. Unpaired two-tailed Student’s t- test was performed to obtain the p values. *** p < 0.001, ** p<0.01, * p<0.05 M- Quantitative PCR-mediated expression studies of *cadA*, *cadB,* and *cadC* genes within infected peritoneal macrophages under the inhibitor treatment of SIRT1 (EX-527) and SIRT3 (3TYP) inhibitors. Data is representative of N=3, n=4. Unpaired two-tailed Student’s t-test was performed to obtain the p values. *** p < 0.001, ** p<0.01, * p<0.05

### Inhibition of SIRT3 triggered mitophagy in the S. Typhimurium-infected macrophages by altering mitochondrial fusion-fission dynamics

Our previous data suggested disruption in mitochondrial bioenergetics in murine macrophages during *S.* Typhimurium infection under SIRT1 and SIRT3 knockdown conditions. To examine whether SIRT1 or SIRT3 knockdown alters mitochondrial dynamics upon *S.* Typhimurium infection, we assessed mitophagy occurrence through immunoblotting and immunofluorescence studies. Using a mito-eGFP-mCherry traffic light construct we observed increased mitophagy induction in SIRT1 and SIRT3 inhibitor-treated infected macrophages at 16hr post-infection (Fig.8A-B). Immunoblotting studies revealed increased mitophagy initiation in the SIRT3 knockdown infected RAW264.7 macrophages as early as 2hr post-infection, with reduced levels of mitochondrial outer membrane protein TOM20 and elevated LC3B II protein levels (Fig. 8C-D). Altogether, our data depicts increased mitophagy influx upon knockdown of SIRT3 in the *Salmonella* infected murine macrophages.

**Fig. 8.**
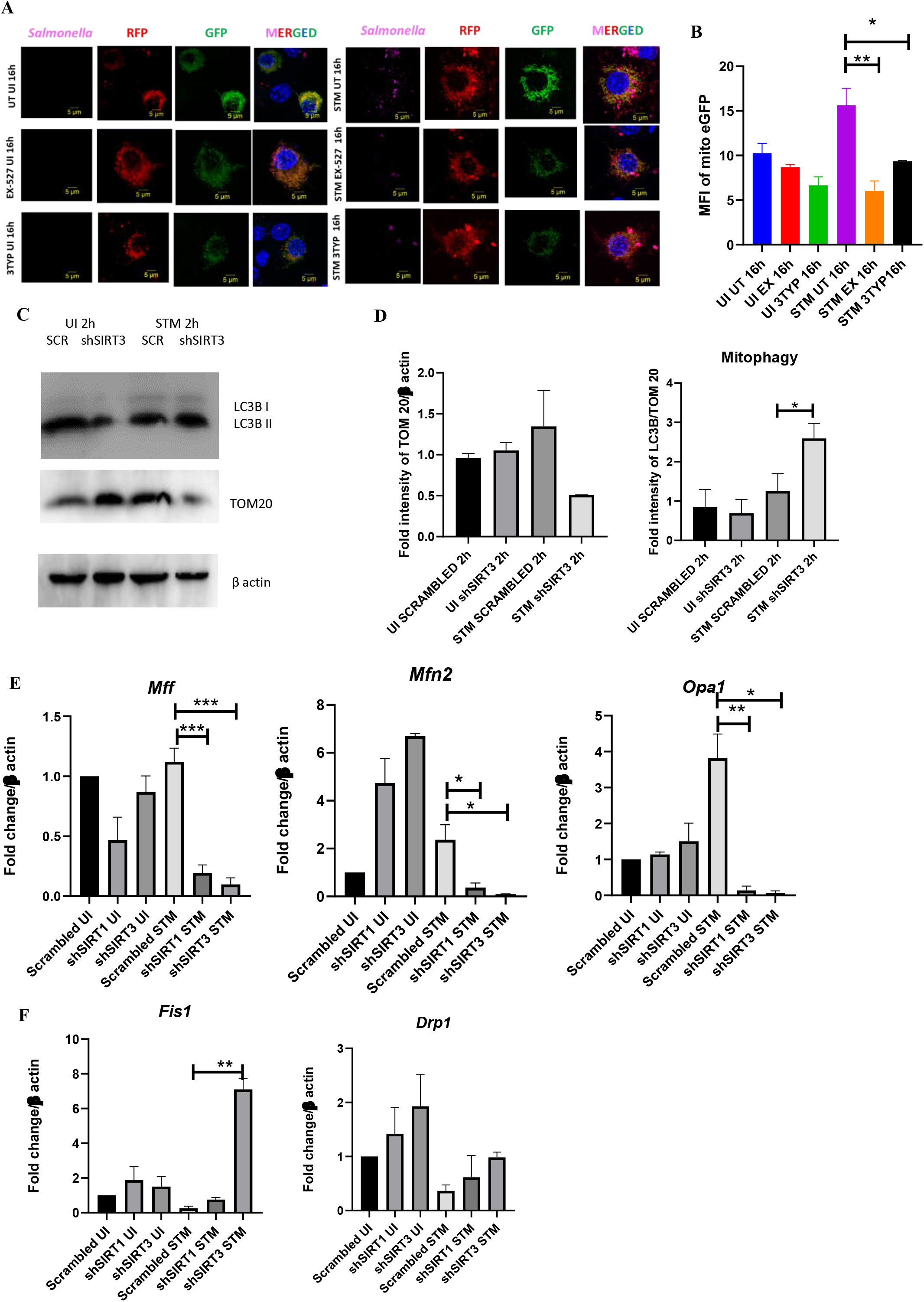
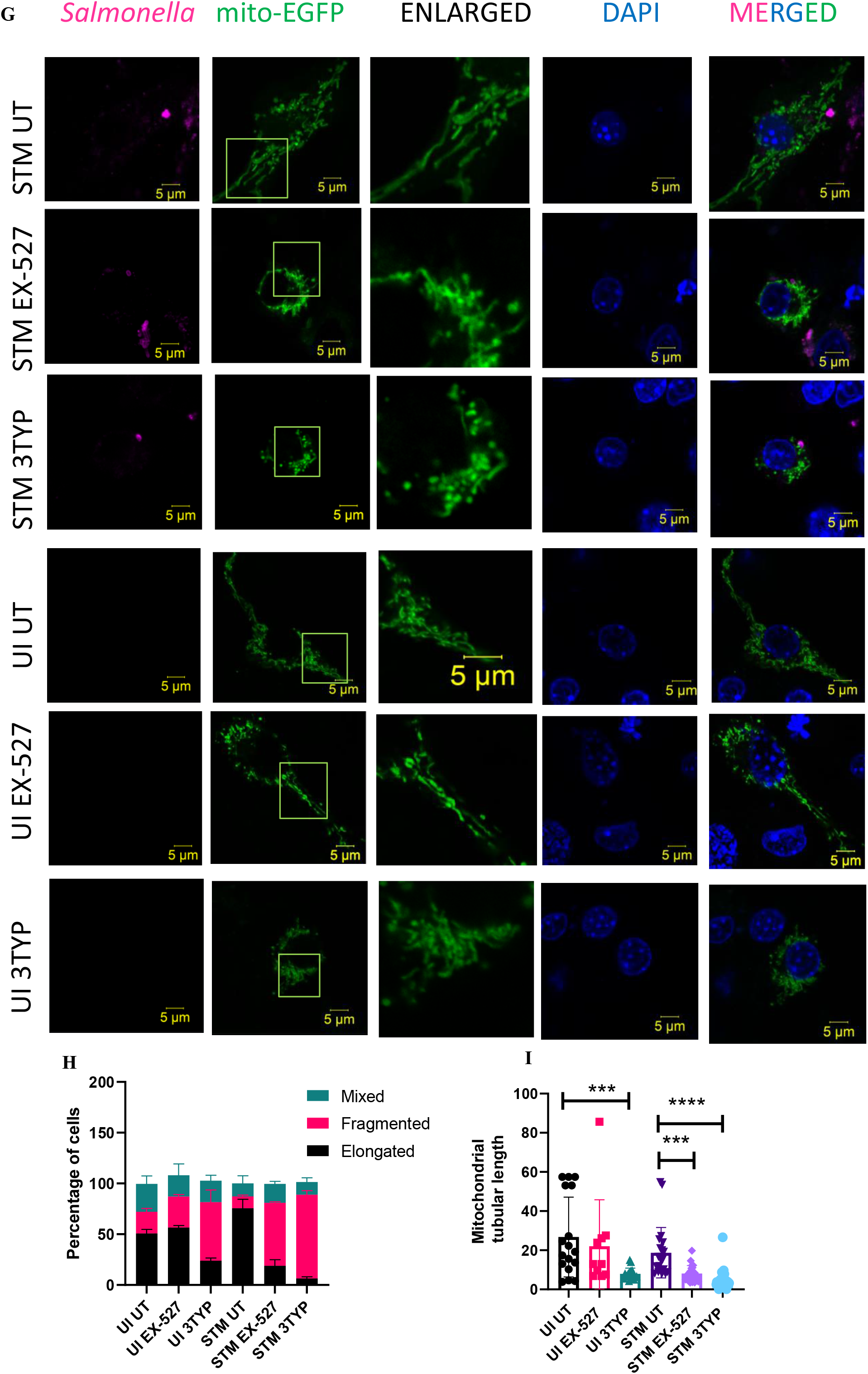
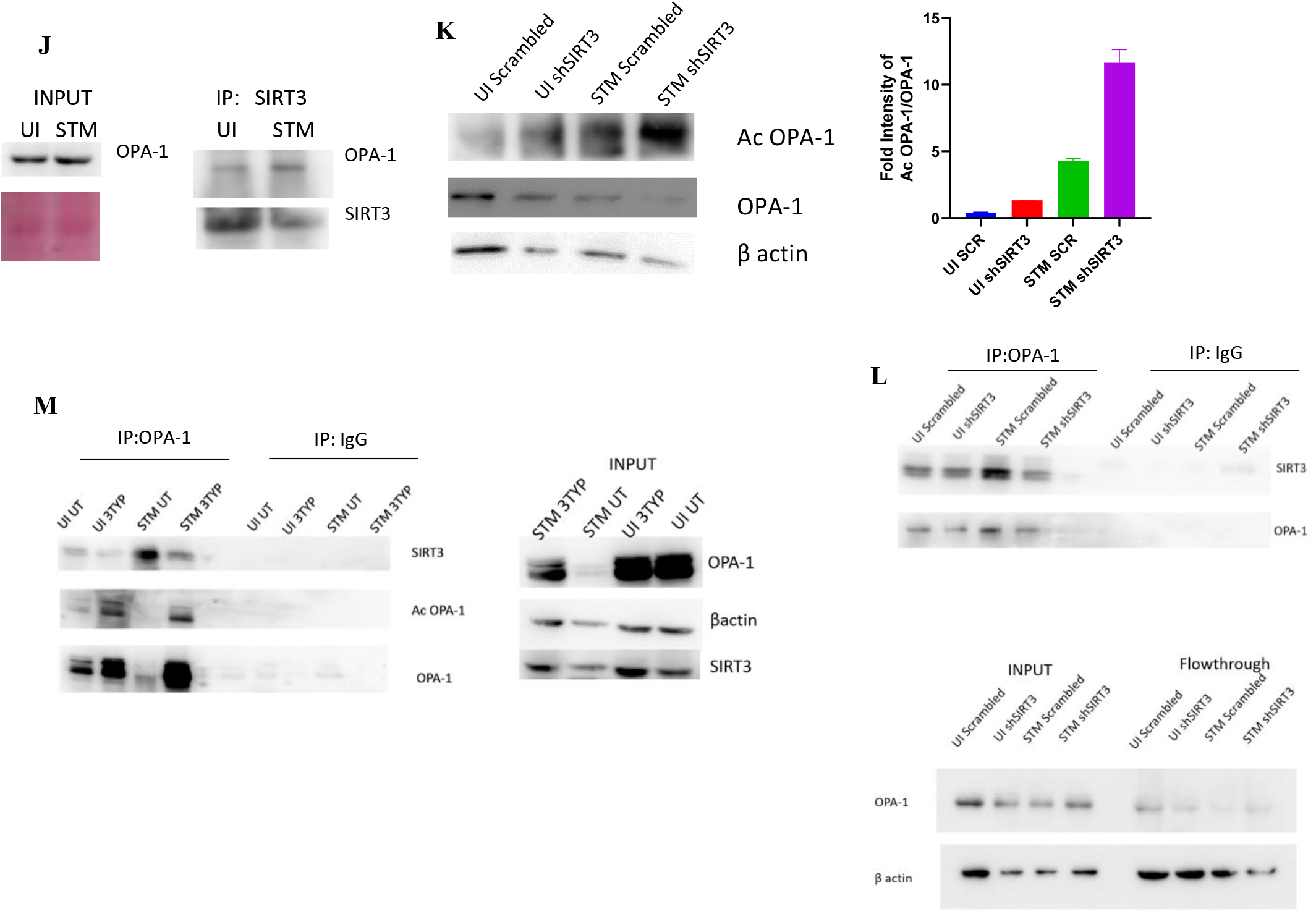
SIRT1 and SIRT3 inhibition triggers mitophagy in the *S.* Typhimurium-infected macrophages by regulating fusion-fission dynamics. A-Representative confocal images of RAW264.7 macrophages (transfected with mito- eGFP-mCherry) exhibiting mitophagy flux upon *S.* Typhimurium infection at 16h points post-infection. Data is representative of N=2, n>50 (microscopic field). B- Quantitative representation of the MFI of mitochondrial eGFP as shown in the confocal images (A). Unpaired two-tailed Student’s t-test was performed to obtain the p values. *** p < 0.001, ** p<0.01, * p<0.05 C- Immunoblotting of LC3B and TOM20 in uninfected and infected RAW264.7 macrophages under knockdown condition of SIRT3 at 2hr post-infection. UI- Uninfected, STM-*S.* Typhimurium infected, Scrambled shRNA control D-Densitometric analysis of TOM20 with respect to β TOM20 and of the immunoblot A to detect mitophagy. E-F- qRTPCR mediated expression analysis of mitochondrial fusion (E) and fission (F) genes in *S.* Typhimurium infected RAW264.7 macrophages at 16hr post infection under knockdown of SIRT1 or SIRT3. G- Representative confocal images of RAW264.7 macrophages demonstrating mitochondrial fusion-fission dynamics upon *S*. Typhimurium infection at 6h post-infection upon SIRT1 or SIRT3 inhibitor treatment. Data is representative of N=2, n>100. H-Quantitative representation of the percentage of cells depicted in (G) exhibiting elongated, fragmented, and mixed population mitochondrial morphologies. I-Quantitative representation of the mitochondrial tubular length of infected (STM) and uninfected (UI) RAW264.7 macrophages as shown in (G). Unpaired two-tailed Student’s t-test was performed to obtain the p values. *** p < 0.001, ** p<0.01, * p<0.05 J- Immunoprecipitation of SIRT3 in uninfected or *S.* Typhimurium infected RAW264.7 macrophages at 16hr post infection to detect interaction of SIRT3 with OPA-1. K- Immunoblotting of SIRT3 knockdown RAW 264.7 cells in *S.* Typhimurium infected and uninfected upon immunoprecipitation of OPA-1 to check the acetylation status of OPA-1. L- Immunoprecipitation of OPA-1 in uninfected or *S.* Typhimurium infected RAW264.7 macrophages at 16hr post infection under the knockdown condition of SIRT3 to assess the interaction of OPA-1 with SIRT3 in knockdown condition. M- Immunoprecipitation of OPA-1 in uninfected or *S.* Typhimurium infected RAW264.7 macrophages at 16hr post infection under the catalytic inhibitor treatment of SIRT3.

Since we observed an increased mitophagy induction upon SIRT3 knockdown condition in infected murine macrophages, we hypothesized whether increased mitophagy coincides with increased incidences of mitochondrial fission. Therefore, we assessed the mitochondrial fission and fusion genes via qPCR in infected SIRT1 or SIRT3 knockdown RAW264.7 macrophages or peritoneal macrophages under the chemical inhibitor treatment of either SIRT1 (EX-527) or SIRT3 (3TYP). qPCR analysis revealed increased transcript level expression of mitochondrial fission genes (*Drp1*, *Fis1*, *Mid49* and *Mid51)* and decreased expression of mitochondrial fusion genes (*Mff*, *Mfn2*, and *Opa1)* in infected macrophages under SIRT1 or SIRT3 knockdown (Fig. 8 E-F) or inhibitor-treated conditions (Fig. EV6) with SIRT3 inhibition showing more prominent effect. Evaluation of mitochondrial dynamics in infected macrophages under inhibitor treatment via immunofluorescence studies revealed an increased percentage of cells exhibiting mitochondrial fusion in *S*. Typhimurium-infected macrophages at 6hr post-infection. However, SIRT1 and SIRT3 inhibitor treatment enhanced mitochondrial fission with around 60% and 80% of cells depicting fragmented morphology, respectively (Fig. 8G) along with reduced mitochondrial tubular length (Fig. 8G). Previous studies have shown the role of SIRT3 in deacetylating OPA-1 and thereby causing activation of OPA-1 in cardiac tissues (Samant et al., 2014). In line with this observation, we evaluated the interaction of SIRT3 with OPA-1 in infected RAW 264.7 macrophages and observed an enhanced interaction in the infected macrophages compared to the uninfected control (Fig. 8 H). However, SIRT3 knockdown or treatment with catalytic inhibitor 3TYP, led to a decline in OPA-1 protein interaction with SIRT3, accompanied by increased OPA-1 acetylation at 16hr post-infection (Fig.8 I-K). These results reveal the role of SIRT3 in controlling the mitochondrial fusion dynamics in *Salmonella-* infected macrophages.

## DISCUSSION

Mitochondrial health emerges as a vital determinant in infection outcomes (Ramond et al., 2019). For instance, pathogens like *L. monocytogenes* and *S. flexneri* induce mitochondrial fragmentation to gain a survival advantage(Stavru et al., 2011, Lum and Morona, 2014) while the intracellular pathogen *Legionella pneumophila* triggers fragmentation via its type IV secretion system effector, MitF, leading to decreased ATP production (Escoll et al., 2017). In contrast, *Chlamydia trachomatis* preserves the mitochondrial framework to maintain the mitochondrial ATP production capacity by preventing Drp-1 mediated mitochondrial fission(Chowdhury et al., 2017) while *C. pneumoniae* causes mitochondrial dysfunction by inducing mitochondrial hyperpolarization, increased ROS production and induction of host metabolic shift to glycolysis(Käding et al., 2017).

Our study investigates how *Salmonella* Typhumurium infection in murine macrophages modulates mitochondrial dynamics and bioenergetics through SIRT1 and SIRT3, influencing infection progression. SIRT3 inhibition increases mitochondrial superoxide generation coinciding with mitochondrial membrane hyperpolarization, proton leakage and decreased ATP production. However, mitochondrial ROS production and membrane hyperpolarization remained unperturbed by inhibition of SIRT1, implicating SIRT3’s predominant role in mitochondrial energy homeostasis and oxidative stress (Ahn et al., 2008, Jing et al., 2011). This aligns with the recent publication on *Mycobacterium tuberculosis* infection wherein SIRT3 deficiency led to increased accumulation of dysfunctional mitochondria with heightened oxidative stress (Kim et al., 2019). While both SIRT1 and SIRT3 deficiencies reduced antioxidant host gene transcript expression, possibly due to their overlapping role in activating the antioxidant defense Nrf2 pathway (Yao et al., 2023, Yang et al., 2020, Kim et al., 2022), mitostress profile showed increased respiration parameters upon both SIRT1 and SIRT3 inhibition. The increased respiratory flux and proton leak gets dissipated in the production of mitochondrial ROS generation leading to decreased ATP content and increased ECAR upon SIRT1 and SIRT3 inhibitor treatment in *S.* Typhimurium infected macrophages. Proton leak and electron slip are the two critical mechanisms contributing to incomplete coupling of ATP generation and substrate oxygen resulting in disproportionate increase in oxygen consumption rate at high protonmotive force and mitochondrial membrane potential with lower ATP yield (Cheng et al., 2017, Kadenbach, 2003, Murphy, 1989).

An important underlying question is how these alterations in the mitochondrial energetics impact the intracellular life of the *Salmonella* within the infected macrophages. Our data indicates that the mitochondrial bioenergetic alteration triggers increased acidification of the macrophage cytosolic pH. SIRT1 or SIRT3 loss or inhibition skews intra-phagosomal and intra-bacterial pH, resulting in decreased SPI-2 gene expression and attenuated intracellular replication.

Numerous evidence suggest how different pathogen assaults lead to remodeling of the mitochondria that eventually dictates the fate of the war between the host and the pathogen (Cervantes-Silva et al., 2021, Carvalho et al., 2020, Shi et al., 2019, Chen et al., 2021, Suzuki et al., 2014). Our work reveals increased mitochondrial fusion dynamics upon *S*. Typhimurium infection via SIRT3-mediated deacetylation of OPA-1. This aligns with *Chlamydia trachomatis* infection studies wherein the infection induces mitochondrial elongation at the early stages and the abrogation of the fusion dynamics led to attenuated chlamydial proliferation(Kurihara et al., 2019). In our study, the *S.* Typhimurium-infected macrophages depicted altered mitochondrial dynamics with increased mitophagy and decreased mitochondrial fusion dynamics under SIRT3 silenced or inhibited condition. The role of mitophagy in host defense varies dictating the infection outcome. During *Pseudomonas aeruginosa* (Kirienko et al., 2015), *Vibrio splendidus* (Sun et al., 2022) or *Mycobacterium tuberculosis* (Lee et al., 2020) infection, host promotes mitophagy for pathogen clearance, while *Listeria monocytogenes* (Zhang et al., 2019), *Yersinia pestis* (Jiao et al., 2022), *Brucella abortus* (Verbeke et al., 2023b) and *Helicobacter pylori* (Jain et al., 2011) (Verbeke et al., 2023a) induce mitophagy for survival or dissemination.

In summary, our results highlight the role of SIRT1 and predominantly, SIRT3 in governing *Salmonella’*s intracellular niche through regulation of mitochondrial energetics, fusion-fission remodeling and host-bacterial intracellular pH balance during *S.* Typhimurium infection. This work contributes to our understanding of host-pathogen crosstalk, offering insights for future avenues in *Salmonella* infection control.

## MATERIALS AND METHODS

### Bacterial Strains, Cell culture condition and infection

*Salmonella* enterica serovar Typhimurium (STM) strain ATCC 14028S, STMΔ*ompR*, STM- pBAD-pHuji or ATCC 14028S constitutively expressing green fluorescent protein (GFP) through pFPV25.1 were used. The above-mentioned live bacterial strain was grown overnight at 37 ^0^C at 160 rpm shaking condition.

| Strains/Plasmids | Characteristics | Source |
| --- | --- | --- |
| *Salmonella enterica* serovar Typhimurium ATCC strain 14028S (STM) | Wild Type (WT) | Gifted by Prof. M. Hensel |
| pFPV:GFP | Amp<sup>R</sup> | Laboratory Stock |
| pFPV:mCherry | Amp<sup>R</sup> | Laboratory Stock |
| pLKO.2 : shRNA | Amp<sup>R</sup> | Gifted by Prof. Subba Rao Gangi Setty |
| STM $\Delta$ *OmpR* | Kan<sup>R</sup> | Laboratory Stock |
| pBAD-pHuji | Amp<sup>R</sup> | Addgene#61555 |
| mito-BFP | Kan<sup>R</sup> | Addgene#49151 |
| STM-pBAD-pHuji | Amp<sup>R</sup> | Laboratory Stock |
| Mito-EGFP-mCherry | Kan<sup>R</sup> | Gifted by Prof. Subba Rao Gangi Setty |

### Cell Culture and Infection Protocol

RAW264.7 macrophages were cultured in DMEM (Lonza) containing 10% Fetal Bovine Serum (FBS) (Gibco) at 37 ^0^C in a 5% CO_2_ incubator. Cells were seeded into 24-well or 6- well plates at 60% confluency.

Peritoneal macrophages from C57BL/6 mice were collected aseptically post-thioglycolate treatment, resuspended in RPMI-1640 (Lonza) with 10% heat-inactivated FBS (Gibco),100 U/ml penicillin and 100 μg/ml streptomycin and seeded into 6 well- plates. 6hr before infection, antibiotic containing media was replaced with Penicillin-Streptomycin-free RPMI- 1640 with 10% FBS.

Macrophages were infected with stationary-phase bacterial culture with MOI of 10. For facilitating the attachment of bacteria to host cells, tissue culture plates were subjected to centrifugation at 600xg for 5 min and plate was incubated at 37 ^0^C humified incubator with 5% CO_2_for 25 min. Cells were washed with PBS and were treated with DMEM (Lonza) + 10% FBS (Gibco) containing 100 μg/ml gentamicin for 1 hr. Subsequently, the gentamicin concentration was reduced to 25 μg/ml and maintained until the cells were harvested for further studies.

### Transfection

PEI-mediated transfection was employed for shRNA-mediated knockdown using plasmids in pLKO.2 vector targeting SIRT1and SIRT3. A control plasmid harbouring scrambled shRNA sequence was also transfected. Plasmid DNA at 500ng per well (24-well plate) was mixed with PEI in 1:2 ratio in serum-free DMEM, incubated for 20 mins and added to RAW 264.7 macrophages. After 6-8hrs, serum-free media was replaced with complete media containing 10% FBS. Transfected cells were either harvested for knockdown confirmation studies or infected with STM 48 hrs post-transfection.

**Table 1:**
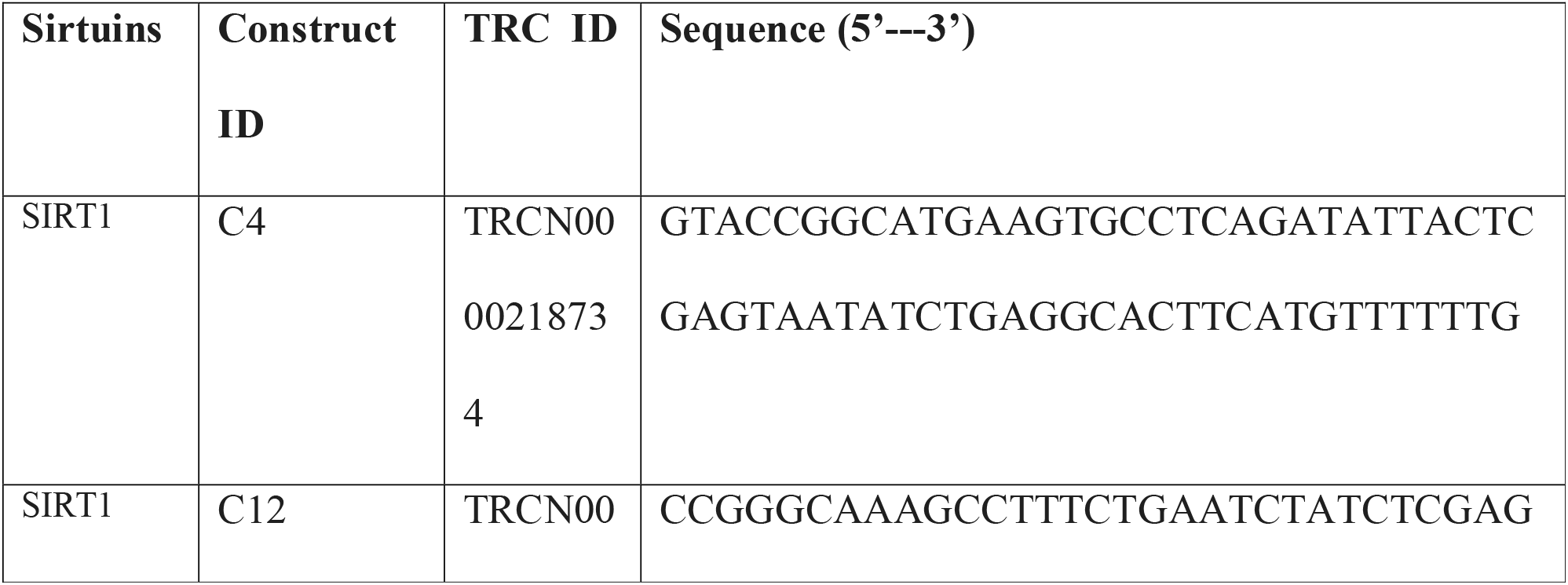
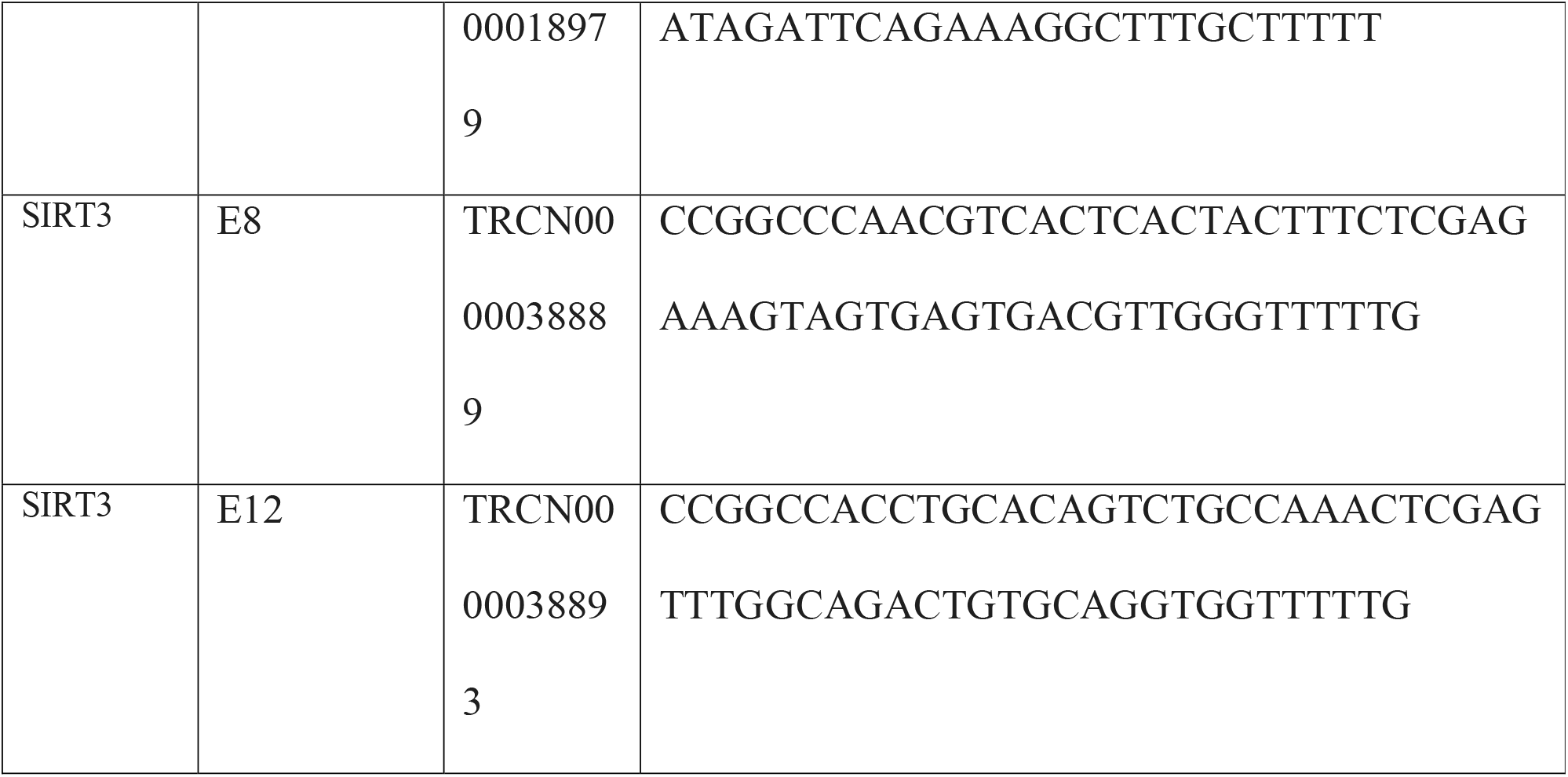
List of shRNA used for knockdown.

### Immunofluorescence confocal microscopic studies

Post-infection with GFP-tagged STM, cells were fixed with 3.5% paraformaldehyde for 15 min at indicated time points. Primary antibody staining, utilizing 0.01% saponin (Sigma) in 2.5% BSA (Bovine serum albumin) containing PBS, occurred overnight at 4°C or for 6hr at room temperature (RT). After PBS wash, cells were stained with appropriate fluorochrome- tagged secondary antibody for 1 hr at RT, followed by DAPI staining and coverslip mounting. Immunofluorescence images were obtained using Zeiss Multiphoton 880 and analyzed using ZEN black 2012 software.

For mitophagy flux and mitochondrial dynamics determination, RAW264.7 macrophages were transfected with mitoGFP-mCherry plasmid. After 24 hrs, RAW264.7 macrophages were infected with *S*. Typhimurium (MOI of 10) and treated with SIRT1 (EX-527) and SIRT3 (3TYP) at 1µM concentration. At designated time points post-infection, cells were fixed, and intracellular *Salmonella* were stained with anti-*Salmonella* antibody. Immunofluorescence images were obtained and analyzed as mentioned earlier. Quantification of mitochondrial tubular length was performed using ImageJ software.

### Quantitative Real Time PCR

Total RNA was isolated at specific timepoints post-infection by using TRIzol (Takara) as per manufacturer’s protocol. RNA quantification was performed in Nano Drop (Thermo-Fischer scientific) and RNA quality was detected via 2% agarose gel electrophoresis. For DNase treatment, 2 µg of RNA was incubated at 37°C for 1 hr, followed by addition of EDTA and heat-inactivation at 65°C for 10 mins. Reverse transcription to cDNA was performed using oligo (dT)_18_ primer, buffer, dNTPs and reverse transcriptase (Takara) as per manufacturer’s protocol. The expression profile of target gene was evaluated using specific primers and SYBR green RT-PCR master mix (Takara) in BioRad Real-time PCR instrument. βactin served as an internal control and all reactions were setup in 384-well plate with two replicates for each sample.

**Table 2:**
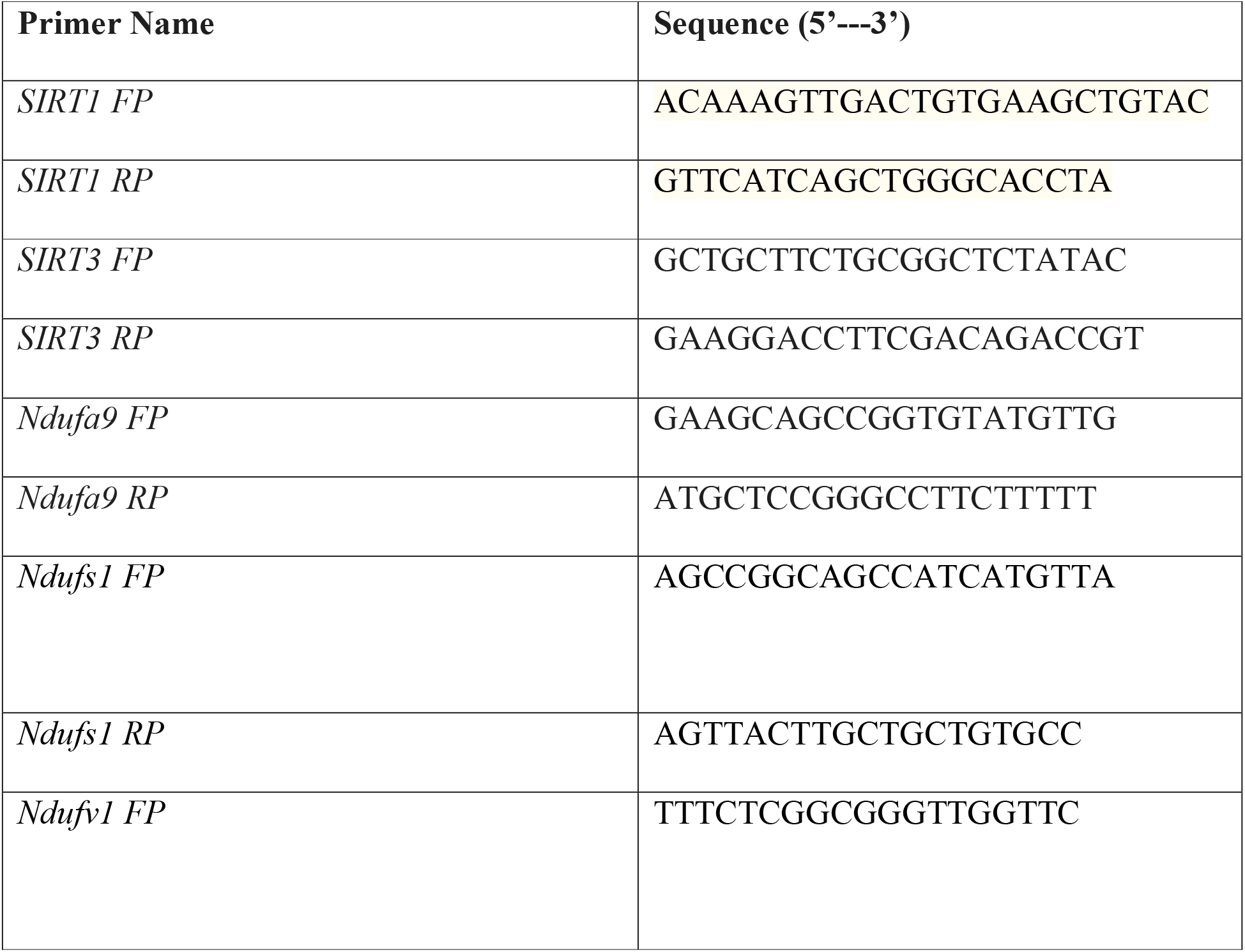
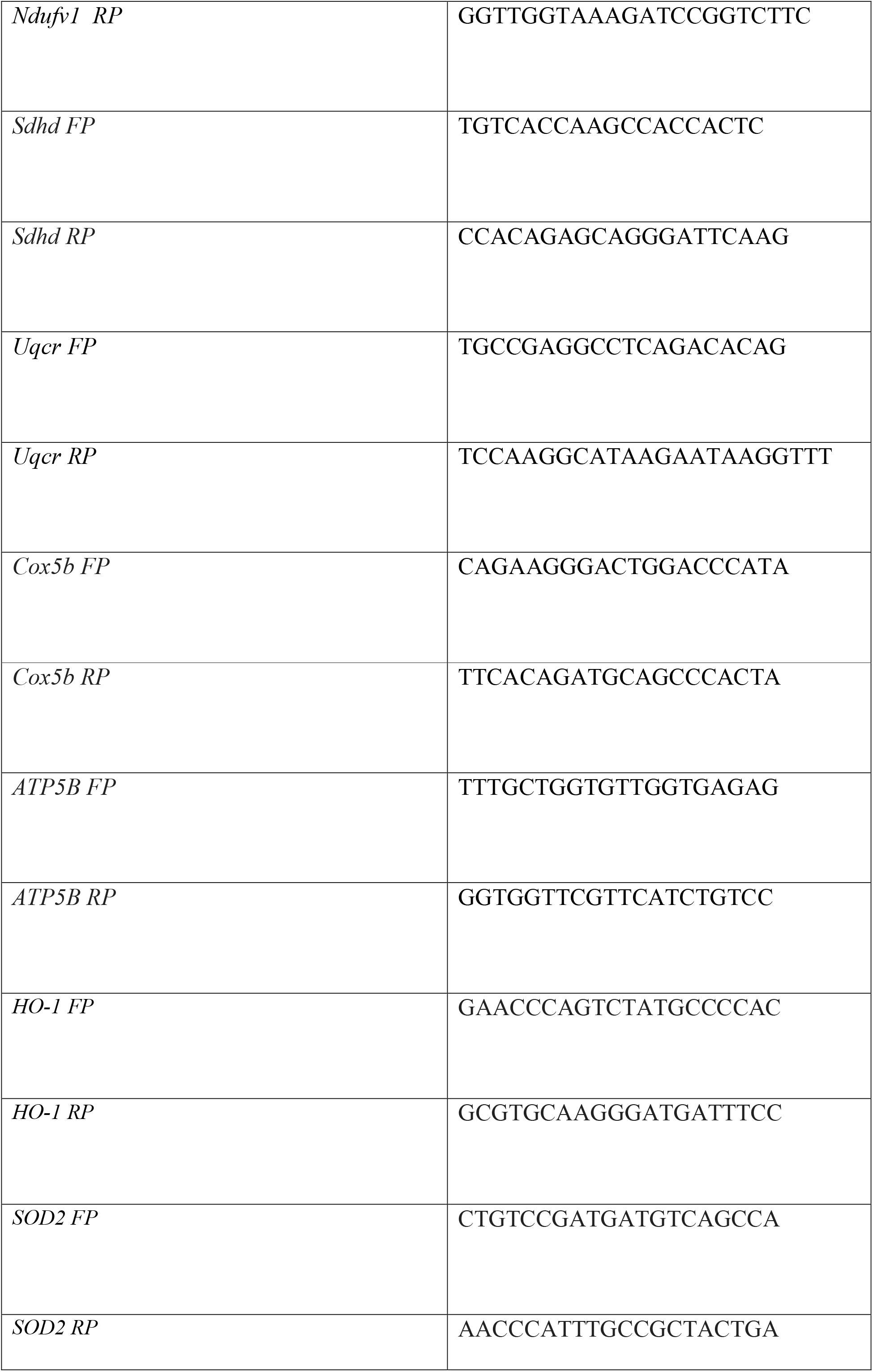
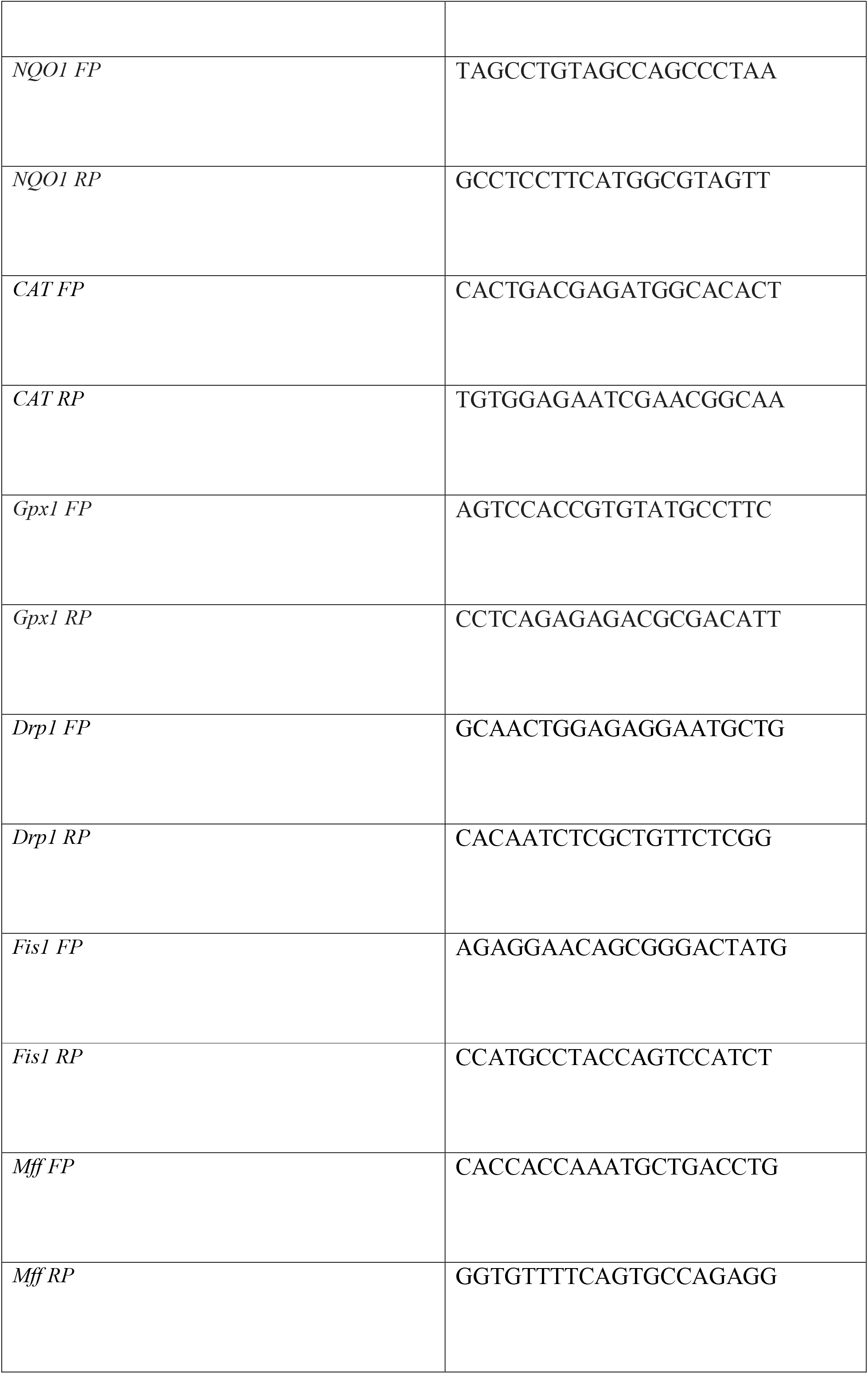
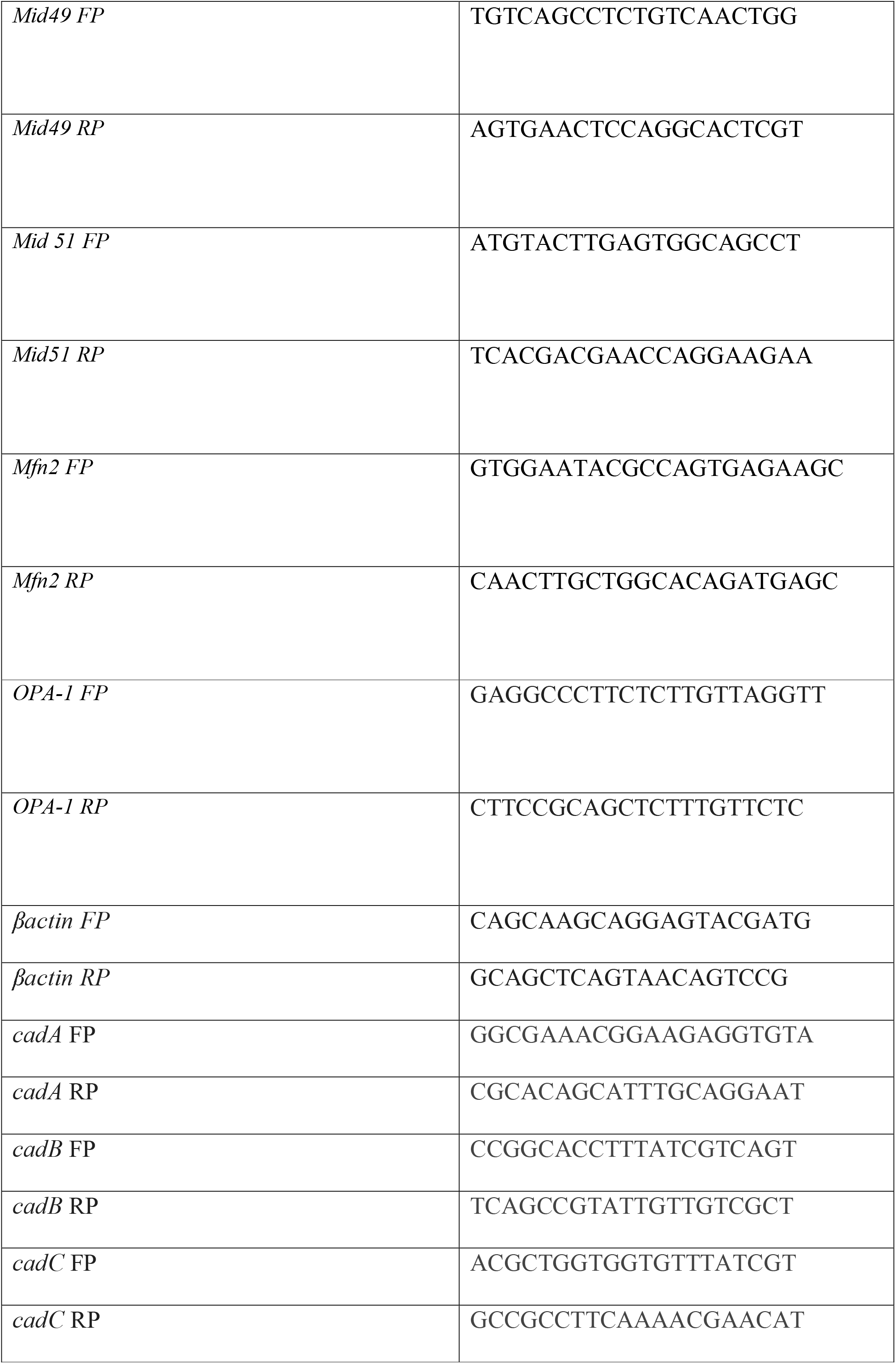
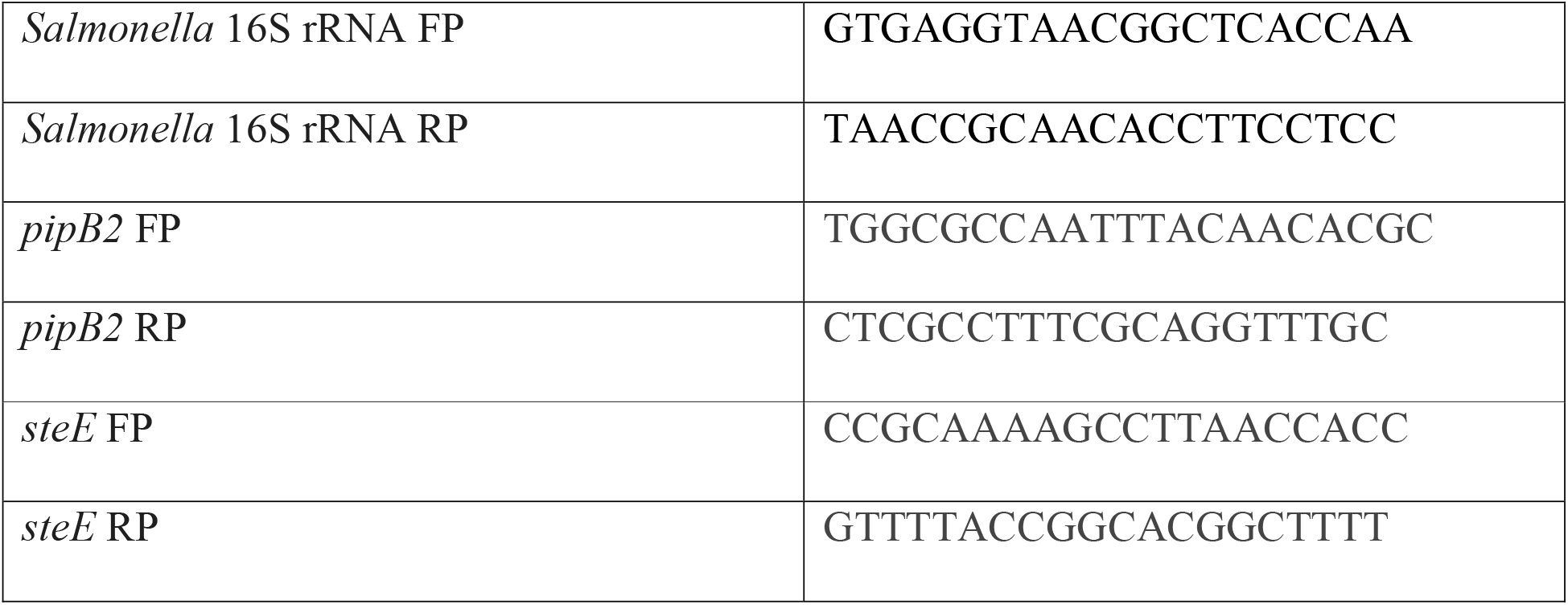
List of primers.

### Intracellular proliferation or gentamicin protection assay

After infecting transfected cells with STM (MOI of 10), they were treated with 100 μg/ml gentamicin containing DMEM (Sigma) + 10% FBS (Gibco) for 1 hr. Subsequently, gentamicin concentration was reduced to 25 μg/ml and maintained until the specified time point. Specific inhibitor treatment was administered with 25 μg/ml gentamicin-containing media at the specified concentration. Post 2hr and 16hr post-infection, cells were lysed in 0.1% triton-X-100, serially diluted and plated to obtain colony-forming units (CFU). Fold proliferation was calculated as CFU at 16hr divided by CFU at 2hr.

### Western Blotting

At appropriate timepoints post-infection, cells were harvested following PBS wash and centrifuged at 1500 rpm for 10 mins at 4°C. Cells were lysed in RIPA buffer containing 10% protease inhibitor cocktail (Roche) for 30 min on ice. Total protein was estimated using Bradford method. Protein samples underwent SDS polyacrylamide gel electrophoresis and were transferred onto 0.45µm PVDF membrane (18V, 2 hrs). The membrane was blocked using 5% skim milk in TBST for 1hr at RT, followed by overnight incubation with appropriate primary antibody at 4°C. After washing in TBST, blot was probed with specific HRP- conjugated secondary antibody for 1hr at RT. The membrane was developed using ECL (Biorad) and images were captured using ChemiDoc GE healthcare. Densitometric analysis was performed using ImageJ software.

**Table 3:**
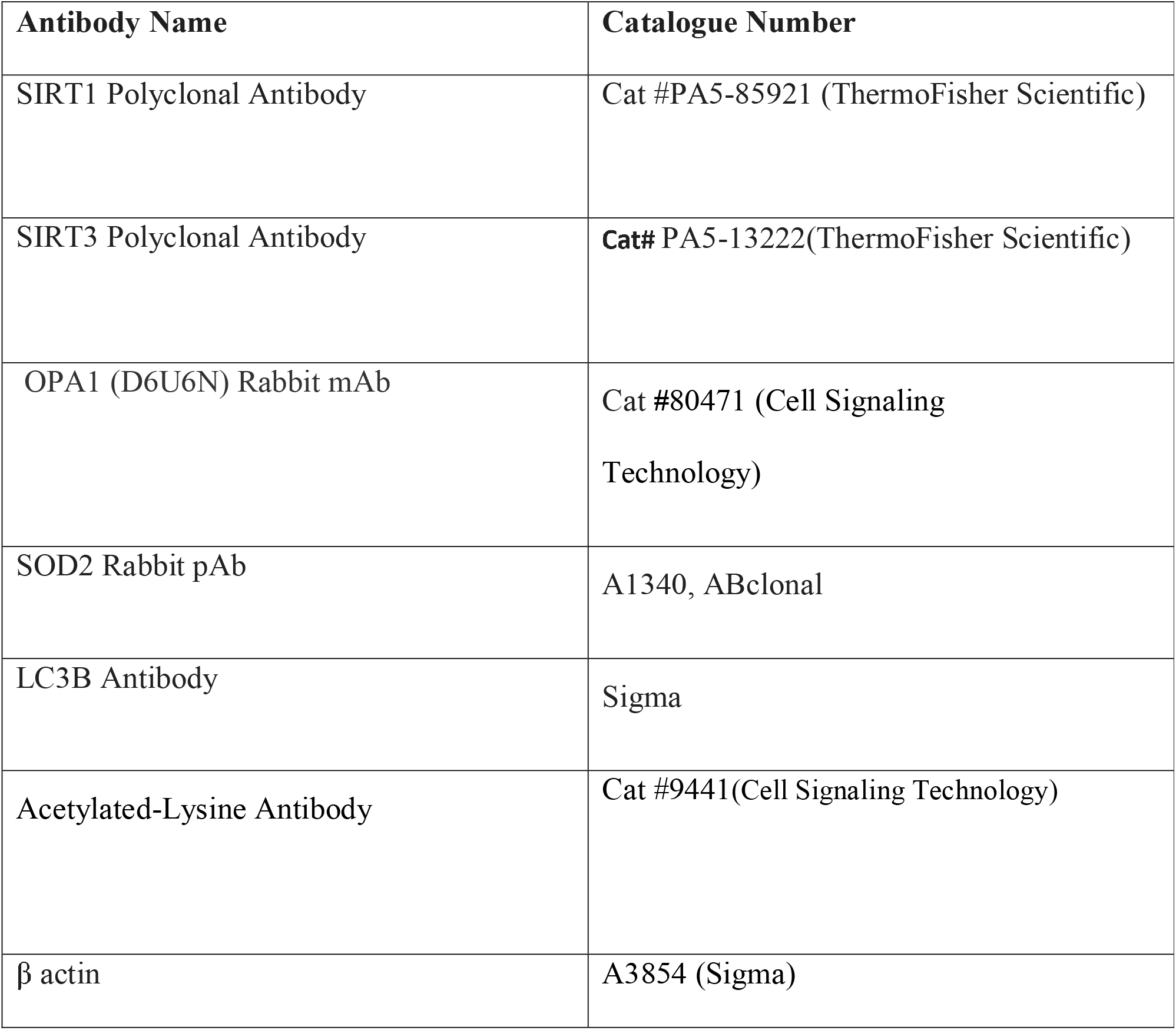
List of Antibodies.

| Antibody Name | Catalogue Number |
| --- | --- |
| SIRT1 Polyclonal Antibody | Cat #PA5-85921 (ThermoFisher Scientific) |
| SIRT3 Polyclonal Antibody | Cat# PA5-13222(ThermoFisher Scientific) |
| OPA1 (D6U6N) Rabbit mAb | Cat #80471 (Cell Signaling Technology) |
| SOD2 Rabbit pAb | A1340, ABclonal |
| LC3B Antibody | Sigma |
| Acetylated-Lysine Antibody | Cat #9441(Cell Signaling Technology) |
| β actin | A3854 (Sigma) |

### Immunoprecipitation

For co-immunoprecipitation, cells were washed with PBS and lysed in native lysis buffer containing 1% Nonidet P-40, 20 mM Tris (pH 8), 2 mM EDTA, 150 mM NaCl and protease inhibitors mixture (Roche Diagnostics) for 30 min at 4°C. Following centrifugation at 10,000 rpm for 10 min, the supernatant was treated with the specific capture antibody. Antibody-lysate complexes were immunoprecipitated using Protein A/G-linked magnetic beads (MagGenome) according to the manufacturer’s protocol. Beads were extensively washed with washing buffer and denatured at 95 °C for 10 min. Denatured precipitates were subjected to the western blotting protocol as mentioned earlier.

### SOD2 Assay

Cell culture supernatant collected at 16h post-infection was analyzed using Mouse Superoxide dismutase [Mn], mitochondrial ELISA kit (SOD2) (ABclonal, Catalog Number- RK04590) following the manufacturer’s instructions. Briefly, 50µl of standards or the test samples were added to each well and incubation was carried for 1 h at 37°C followed by three washes with wash buffer. 100µl of the working Streptavidin-HRP solution was added and incubated for 30 minutes at 37°C. After five washes, 90µL of TMB substrate solution was added per well and incubated for 15-20 minutes at 37°C under darkness. 50 µL of stop solution was added to each well and the optical density was measured within 5 minutes at 450nm with correction wavelengths set at 570nm or 630nm. The concentration of the samples was extrapolated from the standard curve.

### ATP Estimation Assay

Cells harvested at required timepoints post-infection were estimated for ATP content using ATP Colorimetric Assay Kit (Sigma, MAK190) following the manufacturer’s protocol. In duplicates,50µL samples were added per 95-well plate. A master mix containing 44µL of ATP assay buffer, 2µL of ATP probe, 2µL of ATP converter and 2µL of Developer Mix was added. After mixing, the plate was incubated for 30 minutes in dark at RT. Post-incubation, absorbance was measured at 570nm and ATP content was estimated from the ATP standard curve ranging from 0-10 nmole/µl. A treatment of 10µM oligomycin, an ATP synthase inhibitor, was provided to the cells as negative control (Zhou and Faraldo-Gómez, 2018, Shchepina et al., 2002).

### Seahorse OCR Measurement

Mitochondrial oxidative phosphorylation was assessed by measuring the oxygen consumption rate (OCR) using a modified mitostress test(Cumming et al., 2018) on a XF Extracellular Flux Analyzer (Agilent Technologies) at 6hr post-infection following SIRT1/SIRT3 inhibitor treatment. 8-well XF sensor cartridge containing 200 µl/well of XF calibrate solution was preincubated overnight at 37 °C in a CO2-free incubator. 10-fold concentrated compound of oligomycin (Complex V inhibitor), and a combination of rotenone (Complex I inhibitor) and antimycin A (Complex III inhibitor) were loaded into a sensor cartridge to achieve final concentrations of 1.5 µM, and 2 µM, respectively. After a 30-min calibration of the XF sensor with the preincubated sensor cartridge, the cell plate was loaded into the analyzer, and OCR was analyzed under basal conditions. Sequential injection of complex inhibitors was used to assess oxygen utilization for ATP generation and net mitochondrial oxygen consumption. Data were analyzed using WAVE software (Agilent Technologies).

### Flow Cytometry

At specific timepoints post-infection, the cell growth media was removed, and cells were washed with PBS. TMRM (Thermoscientific, cat no. T668) staining solution at 100nM concentration was added to the cells and incubated for 30 mins at 37°C. After washing in PBS, cells were harvested for flow cytometry with 488 nm laser excitation and a 570 ±10 nm emission filter. For MitoSox Red staining, 5µM of MitoSox Red (Invitrogen, cat no.M36008) was used. Cells treated with 10µM of antimycin served as a positive control for mitochondrial ROS production (Han et al., 2008, Park et al., 2007). Post 10-30 mins incubation at 37°C, protected from light, cells were analyzed via flow cytometry. For BCECF-AM staining, cells were stained with 10µM of BCECF-AM (Invitrogen, cat no.B1150) in 1X PBS. Two hours prior to the specific timepoints, cells were treated with 50µg/ml chloroquine in 25µg/ml gentamicin- containing DMEM media (Roy Chowdhury et al., 2022). This chloroquine treated cells served as a control for increased intracellular pH(Di Trani et al., 2007). After incubation for 10-30 mins at 37°C, protected from light, cells were washed before analyzing via flow cytometry (BD FACSVerse) using 405nm and 488nm excitation and 535nm emission channel. The median fluorescence intensity (MFI) of the macrophage population at 488 nm and 405 nm was estimated using BD FACSuite software. The 488/ 405 ratio was determined to estimate the acidification level of cytosol. A standard curve of known pH versus the 488nm/405nm ratio was obtained by exposing the macrophages at different pH buffered solution in presence of 10µM valinomycin and 10µM nigericin. The obtained standard curve was used to interpolate the unknown pH of the test samples.

### Cell Death Estimation Assay

Cell death estimation assay was performed using Propidium iodide (PI) solution (BD Pharmingen, Cat No.-556463) following manufacturer’s protocol by subjecting the cells subjected to PI staining (at concentration of 50µg/ml in PBS (pH 7.4) at the desired time- point post-infection. The samples were analyzed using flow cytometry as described earlier.

### Intra-phagosomal pH measurement

To measure SCV pH of macrophages, a previously reported protocol (Gogoi et al., 2018) was followed. Overnight grown stationary phase cultures of STM WT was labelled with 20 µM pH rhodo red succinimidyl ester (Molecular Probes, Invitrogen) at 37°C for 1h. Bacteria were washed with 0.1 M sodium bicarbonate buffer (pH= 8.4) before and after labelling. RAW264.7 cells were infected with these labelled bacteria at MOI 10. At 6- and 16 h post-infection, macrophages were washed and resuspended in PBS for flow cytometric analysis (CytoFLEX by Beckman Coulter Life Sciences) using 560 nm excitation and 585 nm emission filter. To generate a standard curve of known pH, macrophages were infected with pH rhodo-red-labelled bacteria and subjected to different pH-buffered solution ranging from pH 5.0 to 8.0 in presence of valinomycin (10µM) and nigericin (10µM). Post-equilibration for 1h, macrophages were harvested and subjected to flow cytometric analysis. The percentage pH rhodo-red-positive population was plotted as a function of pH to obtain a standard curve, which was used to obtain the SCV pH.

### Intracellular bacterial pH Measurement

STM WT and STM Δ*ompR* strains, transformed with a pH sensitive plasmid pBAD-pHUji (Plasmid#61555)(Shen et al., 2014), were equilibrated at different pH range (5.0 to 9.0) using a buffered solution with 40µM sodium benzoate. Flow cytometric analysis of the fluorescence intensity ratios produced a standard curve. The above-mentioned strains were used to infect the macrophages (knockdown macrophages or inhibitor treated). 16hr post-infection, the cells were harvested for flow cytometric analysis to obtain fluorescence ratios. The standard curve was used to interpolate ratios measured to obtain intracellular bacterial pH within infected macrophages.

### Statistical analysis

Data were analysed and graphed using the GraphPad Prism 8 software (San Diego, CA). Statistical significance was determined by Student’s t-test or Two-way ANOVA and Bonferroni post-t-test to obtain p values. Adjusted p-values below 0.05 are considered statistically significant. The results are expressed as mean ± SEM.

## AUTHOR CONTRIBUTION

DH and DC have conceptualized the study. DH has contributed to the experiment designing, visualization, methodology, investigation, formal analysis, literature survey, validation, writing (original draft), reviewing and editing of the manuscript. VY has performed the Seahorse mitostress test. AS supervised and provided valuable inputs for the mitochondrial stress test experiments. DC has contributed to the funding acquisition, project administration, and overall supervision of the work. All authors approved the final version of the manuscript.

## DECLARATION OF INTEREST

The authors are unaware of any conflicting interests and thereby declare no conflict of interest.

## ACKNOWLEDGEMENT

This work was supported by the DAE SRC fellowship (DAE00195) and DBT-IISc partnership umbrella program for advanced research in biological sciences and Bioengineering to DC. Infrastructure support from ICMR (Centre for Advanced Study in Molecular Medicine), DST (FIST), and UGC (special assistance) is highly acknowledged. AS acknowledges funding from India Alliance grant IIA/S/16/2/502700), DBT grants (BT/PR13522/COE/34/27/2015, BT/PR29098/Med/29/1324/2018), DST-SERB grant (SPR/2021/000175), Crypto Relief grant (ODAA/INT/20-21), and the Revati and Satya Nadham Atturi Chair Professorship. DH sincerely acknowledges the CSIR- SPM fellowship for her financial support. The funders had no role in study design, data collection and analysis, decision to publish, or preparation of the manuscript. We sincerely thank Prof. Gangi Setty Subba Rao, IISc for providing us with shRNA and mitochondrial constructs.

## EXPANDED VIEW FIGURES

**Fig. EV1.**
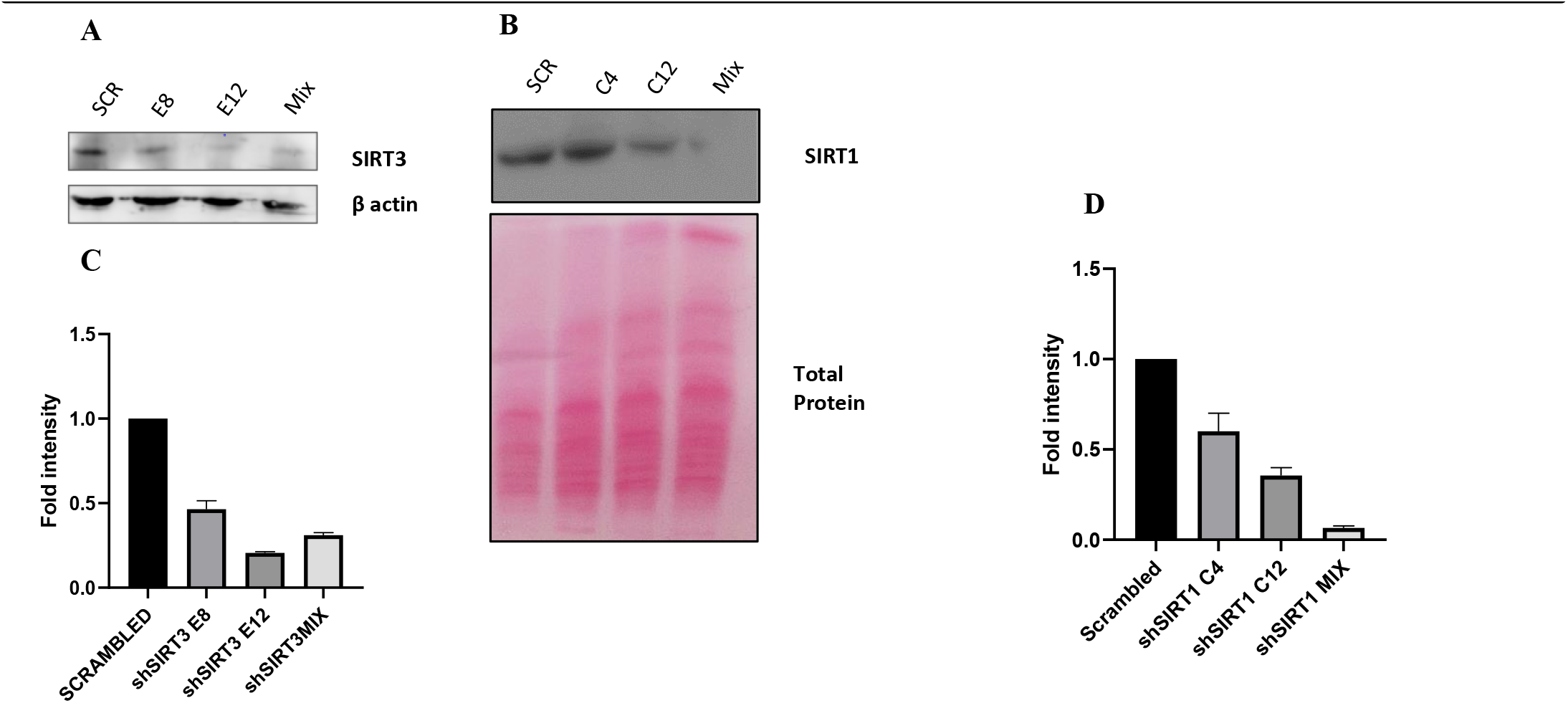
SIRT1 and SIRT3 knockdown confirmation in RAW264.7 cells using shRNA constructs. A- Immunoblotting of SIRT3 for knockdown validation. (SCR=Scrambled, Mix=transfection with equimolar concentration of both SIRT3 shRNA construct E8 and E12) B- Immunoblotting of SIRT1 for knockdown confirmation. (SCR=Scrambled, Mix=transfection with equimolar concentration of both SIRT3 shRNA construct C4 and C12) C- Densitometric analysis of the immunoblot represented in A. D- Densitometric analysis of the immunoblot represented in B.

**Fig. EV2.**
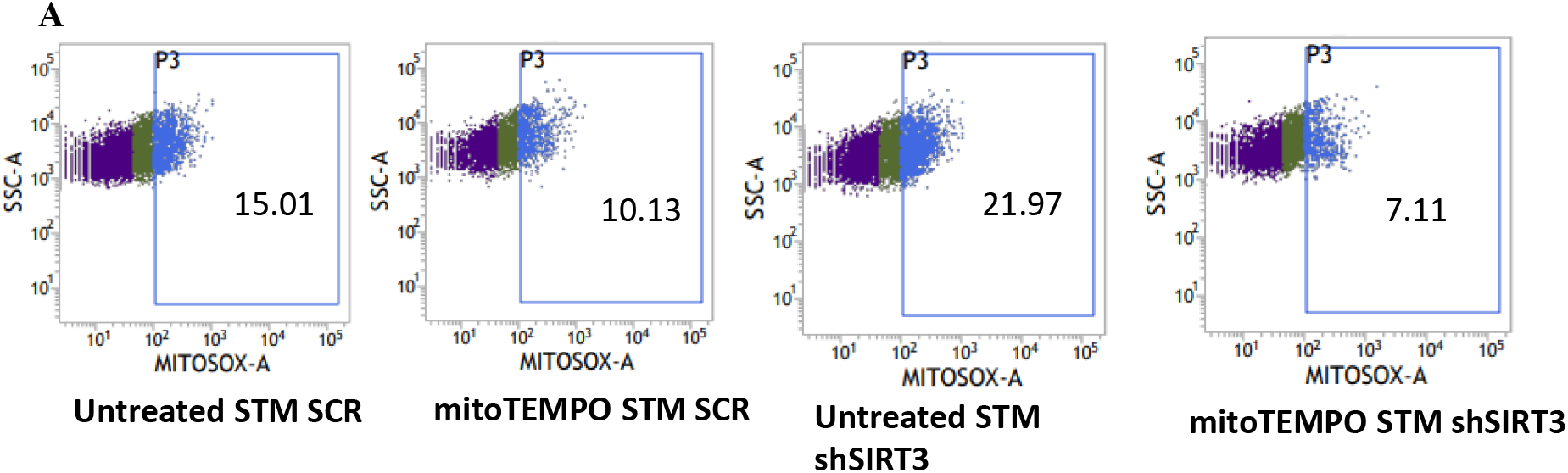
Validation of action of mitoTEMPO as a mitochondrial ROS scavenger. A-Flow cytometric estimation of mitochondrial ROS generation using mitoSOX dye in presence or absence of mtROS scavenger, mitoTEMPO (10µM) in infected RAW264.7 macrophages.

**Fig. EV3.**
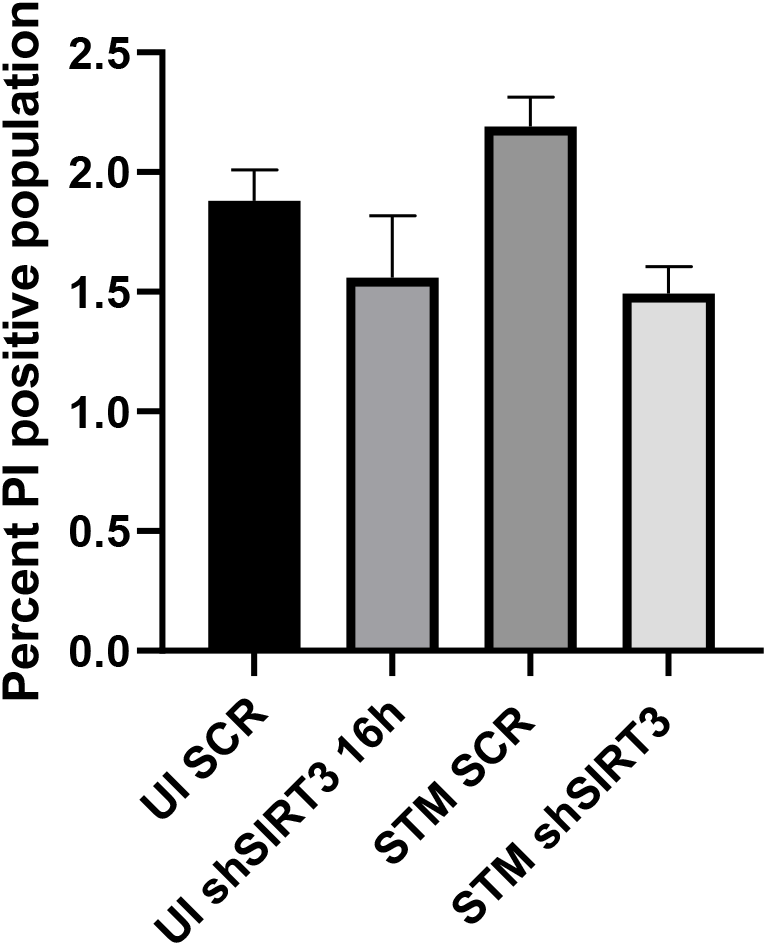
Knockdown of SIRT3 do not impact cell death in the *S*. Typhimurium infected macrophages. Live dead assay via flow cytometric analysis of Propidium Iodide positive RAW264.7 macrophages under knockdown condition of SIRT1 or SIRT3 at 16hr post infection. Data is representative of N=3, n=2. Unpaired two-tailed Student’s t test was performed to obtain the p values.

**Fig. EV4.**
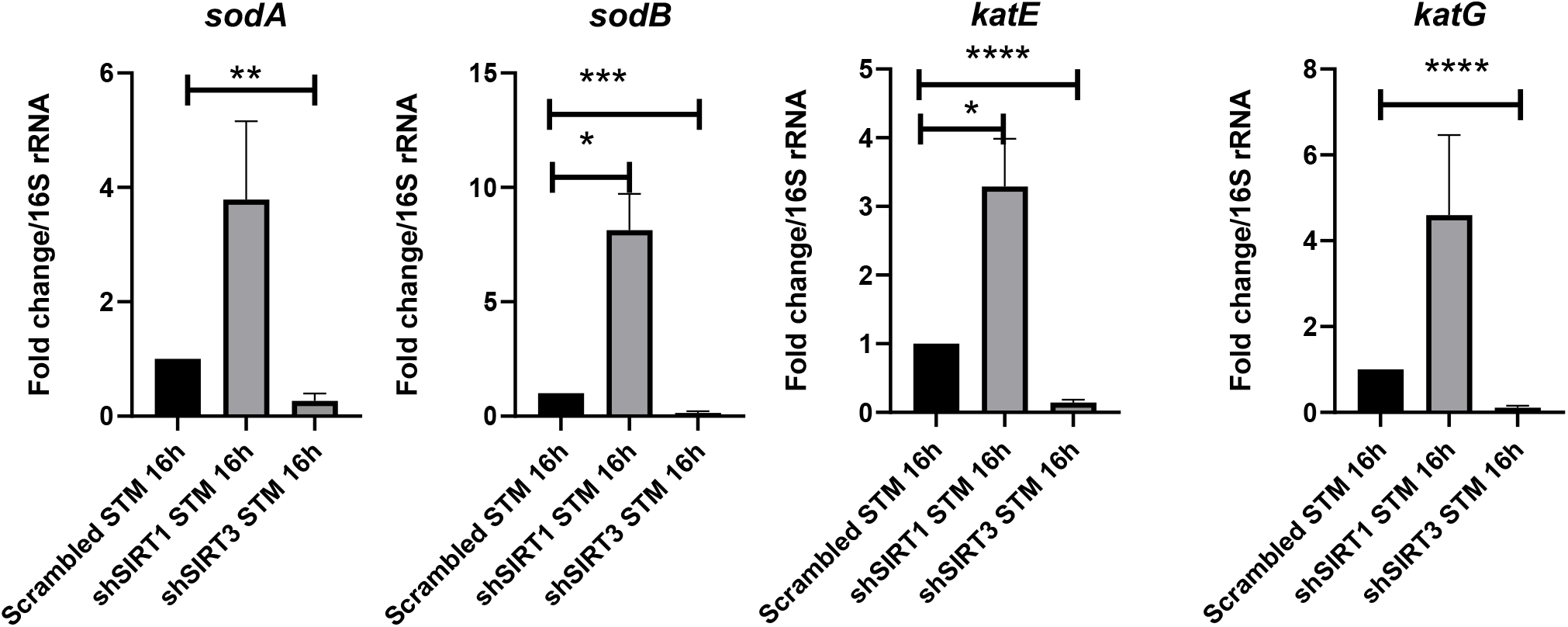
SIRT1 and SIRT3 knockdown lead to alteration in bacterial antioxidant gene expression. qRT-PCR mediated expression analysis of bacterial antioxidant genes in *S.* Typhimurium infected RAW264.7 macrophages at 16hr post infection under knockdown condition of SIRT1 or SIRT3. Data is representative of N=3, n=2. Unpaired two-tailed Student’s t test was performed to obtain the p values. *** p < 0.001, ** p<0.01, * p<0.05

**Fig. EV5.**
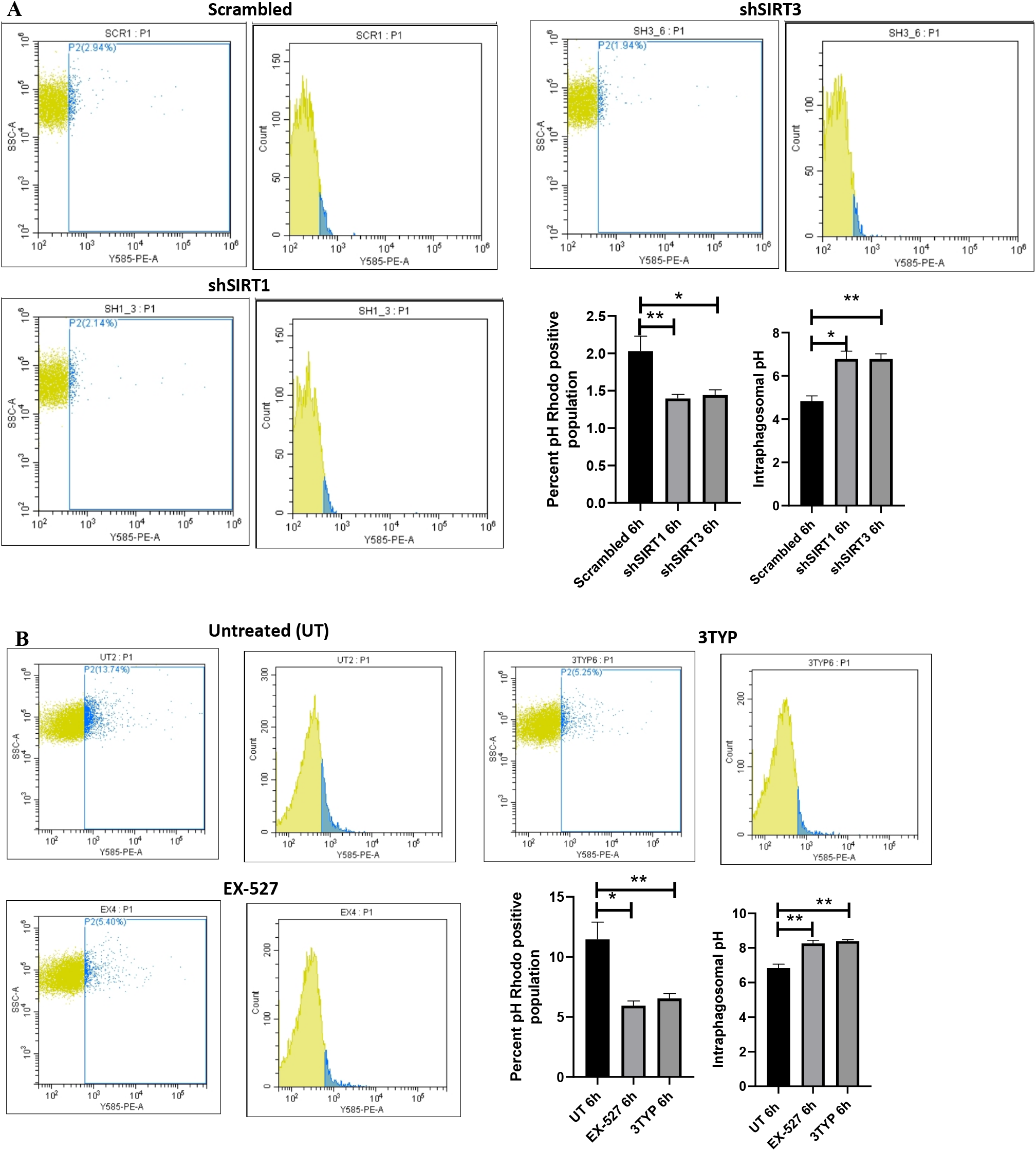
SIRT1 and SIRT3 knockdown or inhibition leads to loss of acidification of SCV. A- Flow cytometric analysis of SCV pH of infected RAW264.7 macrophages under knockdown condition of SIRT1 or SIRT3 at 6h post-infection. Data is representative of N=2, n=3. Unpaired two-tailed Student’s t-test was performed to obtain the p-values. *** p < 0.001, ** p<0.01, * p<0.05 B- Flow cytometric analysis of SCV pH of infected RAW264.7 macrophages under inhibitor treatment of SIRT1 or SIRT3 at 6h post-infection. Data is representative of N=2,n=3. Unpaired two-tailed Student’s t-test was performed to obtain the p-values. *** p < 0.001, ** p<0.01, * p<0.05

**Fig. EV6.**
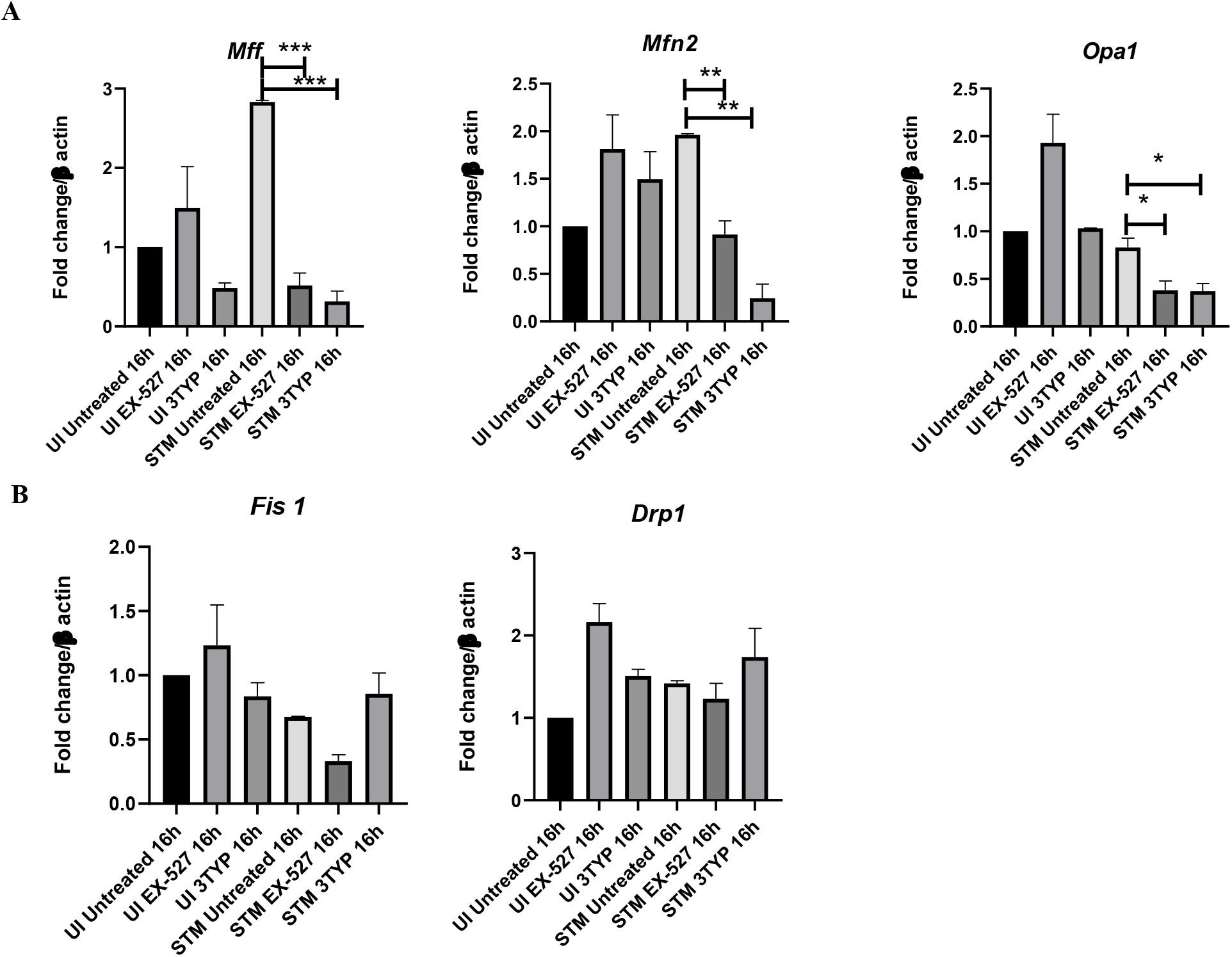
SIRT1 and SIRT3 inhibition causes alteration in mitochondrial fusion-fission dynamics in STM-infected peritoneal macrophages. A- qRTPCR mediated expression analysis of mitochondrial fusion genes in *S.* Typhimurium infected peritoneal macrophages at 16hr post infection under inhibitor treatment of SIRT1 or SIRT3 at a concentration of 1µM. Data is representative of N=3,n=2. Unpaired two-tailed Student’s t test was performed to obtain the p values. *** p < 0.001, ** p<0.01, * p<0.05 B- qRTPCR mediated expression analysis of mitochondrial fission genes in *S.* Typhimurium infected peritoneal macrophages at 16hr post infection under inhibitor treatment of SIRT1 or SIRT3 at a concentration of 1µM. Data is representative of N=3,n=2. Unpaired two-tailed Student’s t test was performed to obtain the p values. *** p < 0.001, ** p<0.01, * p<0.05

## Notes

### Competing Interest Statement

The authors have declared no competing interest.

### Summary of Updates

Figure 8 has been updated with a new data set

